# ATM-deficiency induced microglial activation promotes neurodegeneration in Ataxia-Telangiectasia

**DOI:** 10.1101/2021.09.09.459619

**Authors:** Jenny Lai, Didem Demirbas, Junho Kim, Ailsa M. Jeffries, Allie Tolles, Junseok Park, Thomas W. Chittenden, Patrick G. Buckley, Timothy W. Yu, Michael A. Lodato, Eunjung Alice Lee

## Abstract

While *ATM* loss-of-function has long been identified as the genetic cause of Ataxia Telangiectasia (A-T), how this genetic mutation leads to selective and progressive degeneration of cerebellar Purkinje and granule neurons remains unclear. *ATM* expression is enriched in microglia, the resident immune cell of the central nervous system, throughout cerebellar development and adulthood. Microglial activation has been strongly implicated in neurodegenerative disease and observed in rodent and cellular models of *ATM* deficiency. Here, we find evidence of prominent inflammation of microglia in cerebellum from A-T patients using single-nucleus RNA-sequencing. A-T microglia have transcriptomic signatures of aging and neurodegenerative disease associated microglia. Pseudotime analysis revealed that activation of A-T microglia preceded upregulation of apoptosis related genes in granule and Purkinje neurons, and microglia exhibited increased neurotoxic cytokine signaling to granule and Purkinje neurons in A-T. To confirm these findings experimentally, we studied microglia and neurons that we generated from A-T patient vs. control induced pluripotent stem cells (iPSCs). Transcriptomic profiling of A-T iPSC-derived microglia revealed cell-intrinsic microglial activation of cytokine production and innate immune response pathways compared to controls. Furthermore, adding A-T microglia to co-cultures with either control or A-T iPSC-derived neurons was sufficient to induce cytotoxicity. Taken together, these studies reveal that cell-intrinsic microglial activation may play a critical role in the development and progression of neurodegeneration in Ataxia Telangiectasia.

## Introduction

Selective degeneration of the cerebellum occurs in genetic ataxias such as Ataxia Telangiectasia (A-T). However, the mechanisms underlying cerebellar degeneration remain poorly understood. A-T is an autosomal recessive multi-system disorder caused by mutations in the gene *ATM*^1^. It affects as many as 1 in 40,000 live births and typically presents in early childhood with a median age of diagnosis of 6 years)^2^. Clinical features of A-T include progressive impairment of gait and coordination due to cerebellar neurodegeneration, immunodeficiency, and increased predisposition for cancers. There are currently no therapies to slow neurodegeneration in A-T^2^.

*ATM* is a multifunctional kinase known for its role in DNA damage response to double strand breaks (DSB)^1,3^. DNA damage leads to ATM activation, which phosphorylates downstream regulators of cell cycle arrest, DNA repair, and apoptosis^4^. Additionally, *ATM* plays a role in cellular metabolism. ATM can form a dimer that mediates mitochondrial redox sensing and regulates antioxidant capacity^5^. Moreover, *ATM* deficiency leads to upregulation of autophagy and perinuclear accumulation of lysosomes, implicating *ATM* in lysosomal trafficking^6^.

While *ATM* has roles in diverse cellular functions, it remains unclear how *ATM* loss-of-function leads to selective and progressive degeneration of Purkinje and granule neurons in the cerebellum. ATM-null mice do not recapitulate the cerebellar degeneration found in human A-T patients^7^, but bulk transcriptomic analyses of human brain tissue and neuronal models have begun to offer some insights into dysregulated pathways in A-T. Expression of *ITPR1,* a calcium channel that is highly expressed in Purkinje cells and is associated with spinocerebellar ataxias, is significantly altered in A-T cerebellum^8^. Transcriptome analyses of induced pluripotent stem cell-derived cerebellar-like neurons from individuals with A-T and controls have also revealed alterations in pathways related to synaptic vesicle cycling, oxidative stress, and insulin secretion^9^. However, it remains unresolved whether bulk transcriptomic changes reflect loss of certain cell types in A-T, or perturbation of cellular functions in specific cell types.

Emerging evidence implicates dysfunctional microglia in the pathogenesis of A-T. Human cellular models of microglia with *ATM* deficiency reveal dysfunctional phagocytosis of neuronal processes^10^. However, whether microglia activation is present in A-T patient brains, whether the microglia are reactive to cell-intrinsic or extrinsic signals, and their role in neurodegeneration remain unknown. Characterization of microglia and their transcriptional signatures in human A-T brain may provide further insight on mechanisms that underlie cerebellar degeneration in A-T.

Single-cell technology has provided insight on the cell-type-specific effects of neurological diseases including Alzheimer’s disease (AD), Huntington’s disease, Multiple Sclerosis, Autism, and Major Depressive Disorder^11–15^. However, data from human adult cerebellum in health and degeneration has not yet been explored. Here, we present the largest single-nucleus transcriptomic atlas of adult human cerebellar vermis and the first atlas of cerebellar degeneration to-date. We profiled 126,356 nuclei from the postmortem human cerebellar vermis of six individuals with genetically confirmed A-T and seven matched control individuals. In addition, we profiled 86,354 nuclei from postmortem human prefrontal cortex (PFC) from two A-T and two matched control individuals. We annotated major cell types in the human cerebellum and PFC, and identified cell type proportion changes between A-T and controls. We also demonstrated cell-type-specific expression of cerebellar ataxia-associated disease genes, some of which were significantly dysregulated in A-T Purkinje neurons, supporting a convergence in disease pathophysiology underlying several hereditary ataxias. We analyzed the cell-type-specific molecular pathways perturbed in A-T across these brain regions, revealing prominent and widespread activation of pro-inflammatory pathways in A-T microglia. Pseudotime analysis revealed that activation of A-T microglia preceded upregulation of apoptosis related genes in granule and Purkinje neurons, while ligand-receptor analysis suggested that microglia have increased neurotoxic cytokine signaling to granule and Purkinje neurons in A-T. We experimentally interrogated the role of microglia in neurodegeneration using A-T patient and control iPSC-derived microglia (iMGL) and neurons (iN). Transcriptomic profiling of A-T patient iMGL revealed cell-intrinsic microglial activation of pro-inflammatory pathways, and A-T patient iMGL were sufficient to induce elevated cytotoxicity of co-cultures with control and A-T iNs. Overall, our data suggests that activated microglia are central to A-T pathophysiology and cerebellar degeneration. Our data is publicly available for exploration at the Broad Single Cell Portal (https://singlecell.broadinstitute.org<u>; Study ID SCP130</u>0 and <u>SCP1302)</u>.

## Results

### Single-nucleus RNA-seq profiling of A-T human cerebellum and prefrontal cortex

A-T presents with prominent loss of motor coordination associated with early selective atrophy of the cerebellum. To investigate cell type and region-specific perturbations in A-T, we performed single-nucleus RNA-sequencing (snRNA-seq) of human postmortem tissues from the cerebellar vermis (the region with the most prominent atrophy in A-T^16^) and prefrontal cortex (PFC) (Fig. 1A). We profiled six cerebella from neuropathologically and genetically confirmed A-T cases and seven control cerebella, as well as matched A-T and control PFC from two cases each (Table S1-S4). The ages of the A-T cases ranged from 19 to 50 years old (mean age 28 years). Biallelic ATM variants (with evidence of pathogenicity in ClinVar^21,2217^) were confirmed in all six A-T cases by whole genome sequencing (WGS) (Table S5, Fig. S1). All cases were balanced for age, sex, RNA integrity number (RIN), and postmortem interval (PMI) (Tables S1-S2). After quality control and doublet filtering^18–20^, there were 126,356 cerebellar nuclei (51,297 A-T; 75,059 control) and 86,354 PFC nuclei (16,979 A-T; 69,375 control) for downstream analysis (Fig. S2A-D).

**Figure 1.**
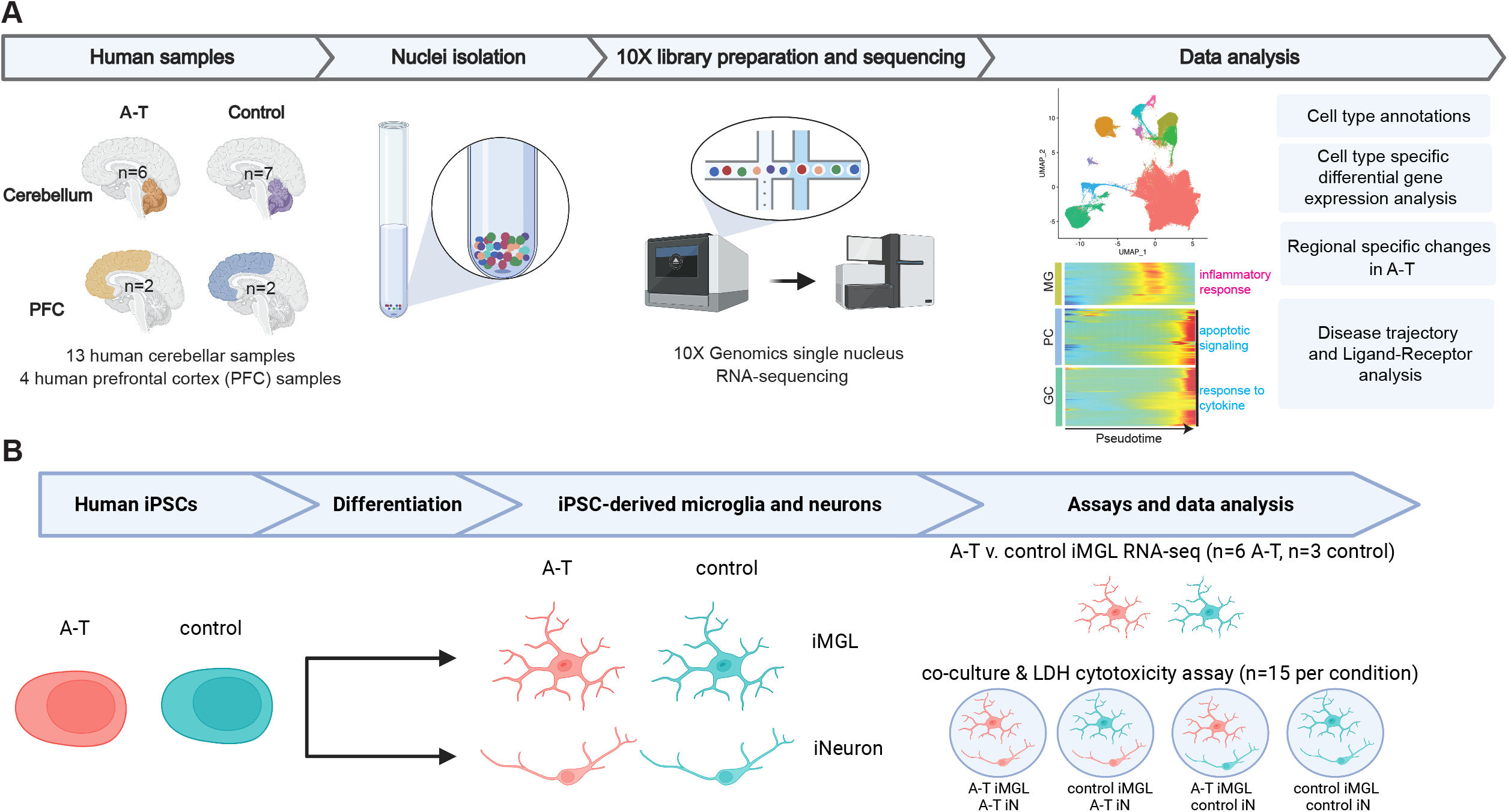
Dissecting cell type specific contributions to neurodegeneration in Ataxia Telangiectasia using single-nucleus RNA-sequencing of postmortem brain and patient iPSC-derived cultures. **A**. Schematic overview of snRNA-seq experimental design and computational analyses. **B**. Schematic overview of A-T patient and control human iPSC-derived experimental design. N represents the number of independent culture wells. MG, microglia. PC, Purkinje cells. GC, granule cells. iMGL, iPSC-derived microglia. iN, iPSC-derived neurons.

### Single-nucleus RNA-seq recapitulates neuropathological hallmarks and reveals increased glial populations in A-T cerebellum

To distinguish cell types in the cerebellum, we used Seurat^18^ to perform graph-based clustering with the Leiden algorithm and annotated clusters based on the expression of canonical marker genes and existing single-cell atlases of the cerebellum^21,22^. Ten major cell types were identified, including the cerebellum specific granule cells (*RIMS1, GRM4)*, Purkinje cells (*ITPR1, CALB1)*, and Bergmann-glia (*TUBB2B, AQP4)*, as well as interneurons (*GAD1, PVALB)*, astrocytes (*TTN, AQP1)*, oligodendrocytes (*PLP1, MBP)*, oligodendrocyte precursor cells (OPC) (*PDGFRA, OLIG1)*, microglia (*CD74, CSF1R)*, endothelial cells (*CLDN5, VWF)*, and fibroblasts (*DCN, APOD)* (Fig. 2A-C). We annotated PFC cell types by reference-based mapping with SingleR, using an existing human PFC snRNA-seq dataset as the reference^13,23^. There were thirteen cell types identified, including astrocytes, microglia, endothelial cells, oligodendrocytes, OPCs, and multiple subtypes of excitatory and inhibitory neurons (Fig. S3A).

**Figure 2.**
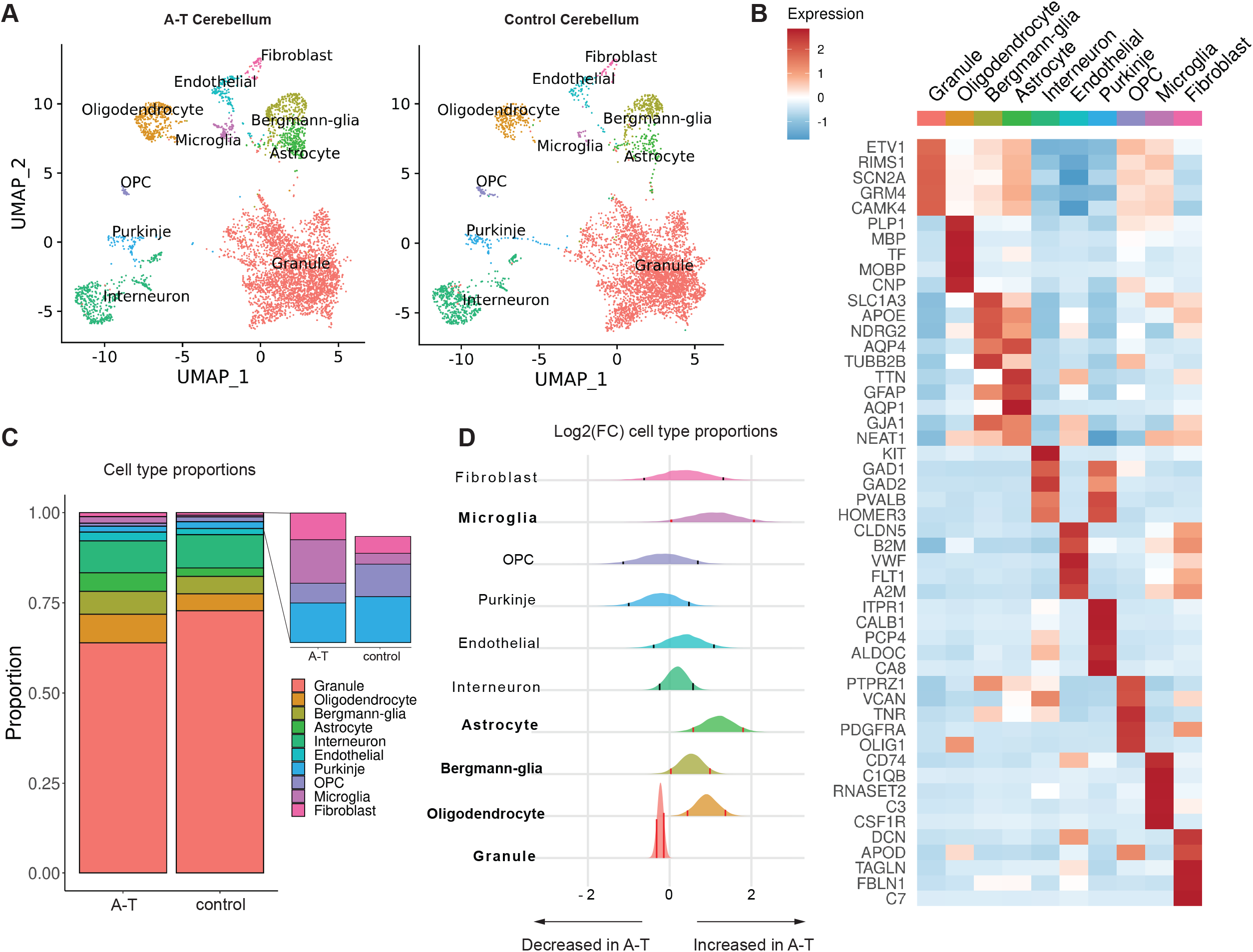
Cell types resolved in A-T and control human cerebellum by snRNA-seq. **A**. UMAP dimensionality reduction plot of major cell types identified in A-T and control human cerebellum, downsampled to 10,000 cells for each condition. **B.** Top five marker genes for each major cell type in control cerebellum. Heatmap depicts centered and scaled log-normalized expression values. **C**. Cell type proportions in A-T and control cerebellum. **D**. Relative abundance of cell types in AT versus control cerebellum, shown as the posterior distribution of Log2(proportion in AT/proportion in control) with 89% credible interval. Red bars highlight credible intervals that do not overlap 0. Bolded cell type labels indicate a significant difference in relative abundance.

Next, to understand the patterns of cerebellar cell loss in A-T, we examined the cell type proportions in A-T compared to controls. Due to the negative covariance structure of compositional data—as one feature increases, the other features must decrease—we implemented Dirichlet multinomial modeling using Hamiltonian Monte Carlo (DMM-HMC) to estimate the cell type proportions in A-T and control^24^. We generated multinomial distributions of cell type proportions for each sample based on the nuclei counts (Table S3), and used these to construct a Dirichlet distribution of cell type proportions in A-T and controls. To determine the relative shift in abundance of each cell type in A-T versus control, we subtracted the control posterior probability distribution of proportions from the A-T distribution (Fig. 2D). Granule cells were significantly decreased in A-T (Fig. 2D, 89% credibility interval). Purkinje cells were reduced in abundance, although not significantly, possibly due to individual sample variability and decreased power for detecting changes in rare cell types. In addition, astrocytes, Bergmann-glia, microglia, and oligodendrocytes showed significant increases in A-T compared to control (Fig. 2D, 89% credibility interval). In contrast, permutation testing of disease labels (A-T versus control) showed no significant differences for any cell type, as expected (permutation n=500, Fig. S2E). Unlike A-T cerebellum, there was no evidence of neuronal loss in A-T PFC (Fig. S3B). However, oligodendrocytes were significantly decreased in A-T PFC compared to control (Fig. S3B), consistent with reports of cortical white matter degeneration in A-T^25,26^. Taken together, these results are consistent with the neuropathology autopsy reports from individuals with A-T, which describe granule and Purkinje cell atrophy and gliosis in the cerebellum while the PFC is grossly unremarkable (Table S4).

### Monogenic cerebellar disease genes are expressed in specific cell types

Next, we characterized the expression of *ATM* across cell types in the cerebellum and PFC. *ATM* had the highest expression in microglia in adult control human cerebellum and PFC (Fig. S4A). To investigate human developmental expression patterns, we profiled the expression of *ATM* in a snRNA-seq dataset of human control fetal cerebellum^27^, which also revealed the highest expression of *ATM* in microglia (Fig. S4B). Taken together, these data suggest an important role for *ATM* in microglia during normal cerebellar development.

Given that *ATM* is enriched in microglia, we asked whether additional hereditary ataxia genes are enriched in specific cerebellar cell types. We curated a list of genes implicated in human hereditary ataxias using the Online Mendelian Inheritance in Man (OMIM) database (Table S6) and profiled their expression in control cells. We found that like *ATM*, most hereditary ataxia disease genes demonstrate cell-type-specific patterns of expression (Fig. S4C; Bonferroni adjusted p-value < 0.05, t-test). Cerebellar disease gene expression was most commonly enriched in Purkinje cells, including the potassium and calcium ion related genes *KCND3, PRKCG, CACNA1G, ITPR1, TRPC3,* and *KCNC3*. Microglia also showed enriched expression of several ataxia disease genes (*ATM, TPP1, SETX,* and *VPS13D*) (Fig. S4C). These gene expression patterns implicate Purkinje cells and microglia in the pathogenesis of cerebellar ataxias.

### Transcriptional dysregulation across cell types in A-T

To investigate the cell types and molecular pathways that are most perturbed in A-T cerebellum, we performed differential gene expression analysis between A-T and control for each cell type (Table S7). Astrocytes, microglia, and oligodendrocytes had remarkably high numbers of DEGs (n=1,631, 1,577, 1,563 respectively) while granule cells had the least (n=265). These data suggest that while granule cells degenerate in A-T, their gene expression program is less dysregulated than other cerebellar cell types such as glia.

Next, we performed Gene Ontology (GO) analysis on the DEGs found in each cerebellar cell type to gain insight into the biological processes dysregulated in A-T. Consistent with the loss of Purkinje and granule cells in A-T, both cell types had significant upregulation of genes involved in apoptotic signaling (Fig. 3A-B, Fig. S5A, Table S8). A-T Purkinje and granule cells also upregulated genes involved in ribosome assembly and biogenesis, which has been observed in several studies of aging brains^28,29^. A-T microglia specifically demonstrated upregulation of immune response related genes involved in microglial cell activation, phagocytosis, and cytokine production (Fig. 3C).

**Figure 3.**
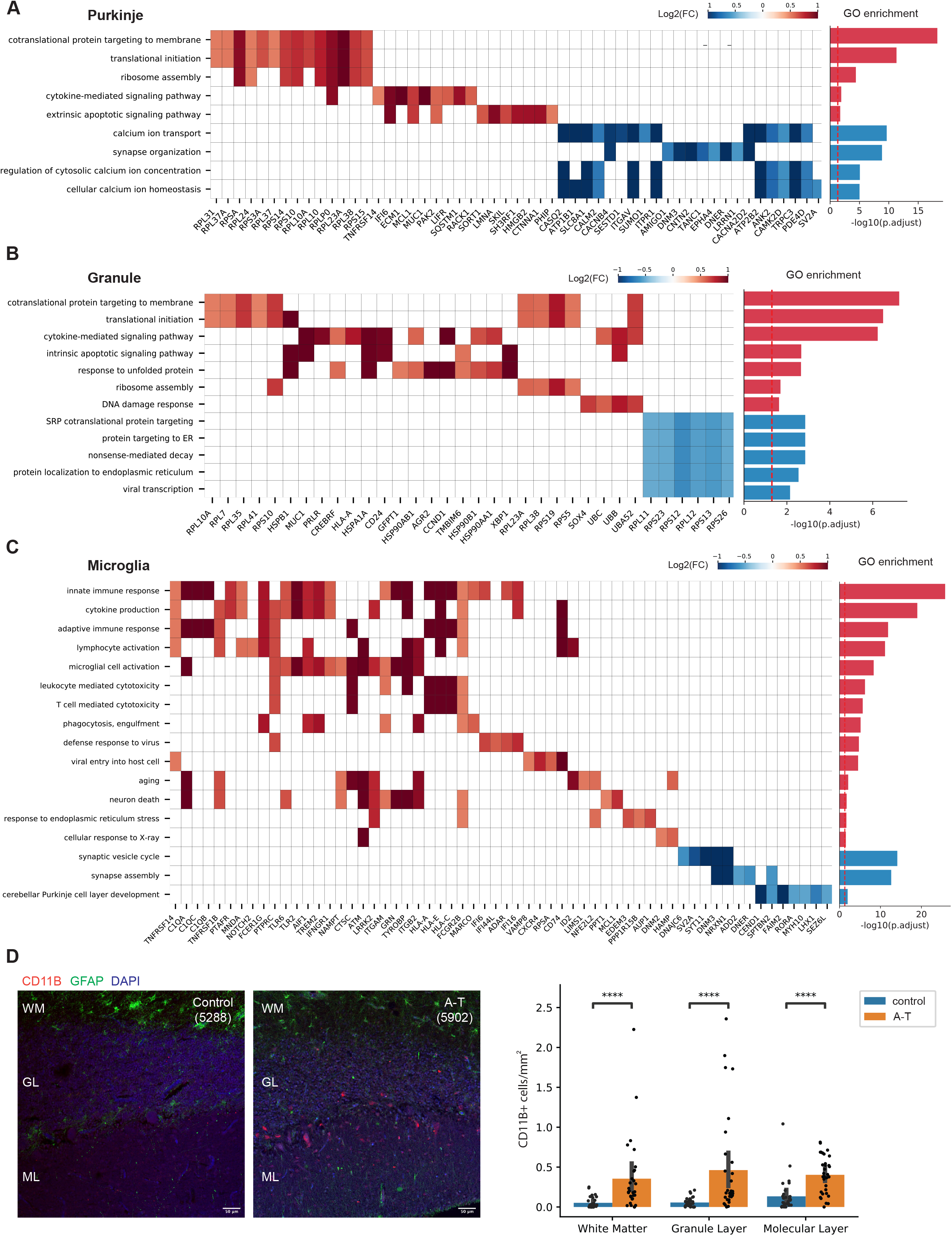
Gene Ontology (GO) analysis of differentially expressed genes in A-T cerebellum. **A-C**. Select enriched GO biological processes among DEGs in A) A-T Purkinje neurons, B) granule neurons, C) microglia. Select genes in each pathway and their log2 fold-changes (A-T/control) shown in the heatmap. Barplot shows significance of pathway enrichment among upregulated genes (red) and downregulated genes (blue). **D**. CD11B immunostaining of postmortem cerebellar cortex from A-T and control. WM: white matter. GL: granule layer. ML: molecular layer. Quantification of CD11B-positive cells across cerebellar cortex layers in AT and control. Error bar represents 95% confidence interval. ****Bonferroni-adjusted p-value<1e-04, Mann-Whitney U test.

Activated microglia associate with sites of injury and pathological changes across various neurodegenerative diseases^30^ and can induce local neurodegeneration by phagocytosis of synapses and secretion of neurotoxic cytokines^31^. We thus sought to characterize the spatial distribution and number of microglia in A-T and control cerebellum by immunostaining for CD11B, a microglia marker that is upregulated in activated microglia^32^, in four A-T cerebellum and three matched control cerebellum samples (Table S9). In A-T cerebellum, there were increased CD11B+ cells throughout the white matter, granule layer, and molecular layer compared to control cerebellum (Fig. 3D). Across all layers, the number of CD11B+ microglia per area was significantly higher in A-T compared to control (Bonferroni adjusted p-values< 1e-04, Mann-Whitney U test) (Fig. 3D). These data confirm that A-T cerebellum has a widespread increased presence of activated microglia.

### Dysregulated calcium signaling in Purkinje neurons in A-T

GO analysis of genes downregulated in A-T revealed significant enrichment of calcium related pathways (calcium ion transport, regulation of cytosolic calcium ion concentration, cellular calcium ion homeostasis) in A-T Purkinje cells (Fig. 3A, Fig. S5, Table S10). Among these genes was *ITPR1,* an endoplasmic reticulum (ER) calcium channel that causes spinocerebellar ataxias in an autosomal dominant manner^33,34^ (Fig. 3A). We asked whether additional cerebellar disease genes are differentially expressed in A-T. Hereditary ataxia genes with enriched expression in Purkinje cells (i.e., *KCNC3, PRKCG, ITPR1, KCND3, CACNA1G)* were significantly downregulated in A-T Purkinje cells (Fig. S7A). These disease genes were also enriched for calcium processes (calcium ion transport and regulation of cytosolic calcium ion concentration) (Fig. S7B). These results suggest that *ATM* may be an upstream activator of calcium signaling and points to converging underlying mechanisms of a subset of hereditary ataxias in Purkinje cells. *HDAC4* is a known negative regulator of calcium ion genes (*ITPR1, DAB1)*^35^, and *ATM* is an inhibitor of *HDAC4*. Loss of *ATM* leads to nuclear accumulation of HDAC4, which could lead to dysregulated expression of *HDAC4* target genes^36^. In support of this, there was significant overlap between *HDAC4* targets and A-T downregulated genes (p-value 6.08e-22, Fisher’s exact test, Fig. S7C). Taken together, these findings support that abnormal calcium ion homeostasis may be a consequence of *ATM* loss of function in Purkinje neurons, which may lead to aberrant firing patterns and ataxic phenotypes^37,38^.

### Activated microglia in A-T cerebellum share transcriptomic signatures with aging and neurodegeneration-associated microglia

Microglia in A-T cerebellum express activation markers, including strong upregulation of the complement genes, *C1QA, C1QB, C1QC,* and *C3* (Fig. 4A). Increased expression of complement is associated with microglial activation in neurodegenerative diseases such as Alzheimer’s disease (AD)^39^, where they have been shown to mediate synaptic loss and dysfunction^40,41^. To assess whether A-T cerebellar microglia share additional features associated with microglia in aging and other neurodegenerative disorders, we compared the upregulated genes in A-T cerebellar microglia with three gene sets: markers of aged human microglia, AD associated microglia markers from a human snRNA-seq study of AD, and markers of disease associated microglia (DAM) from a murine model of AD^11,42,43^. We found significant overlaps between A-T cerebellar microglia upregulated genes and these three gene sets with the highest overlap with aging microglia (Fisher’s exact test p-values < 1e-13; Fig. 4B-C). *TREM2, TSPO, MS4A6A,* complement components, and MHC class II genes were among the overlapping genes, which have been associated with neuroinflammation and microglia activation^46^ (Fig. 4C). GO enrichment analysis of the overlapping genes and A-T microglia specific upregulated genes revealed common enrichment in immune response and RNA processing related pathways, and A-T specific enrichment of cytoskeletal organization, ER stress, and regulation of neuron projection development (Fig. 4D). Overall, these data show that A-T cerebellar microglia share transcriptomic signatures found in aging and other neurodegenerative diseases.

**Figure 4.**
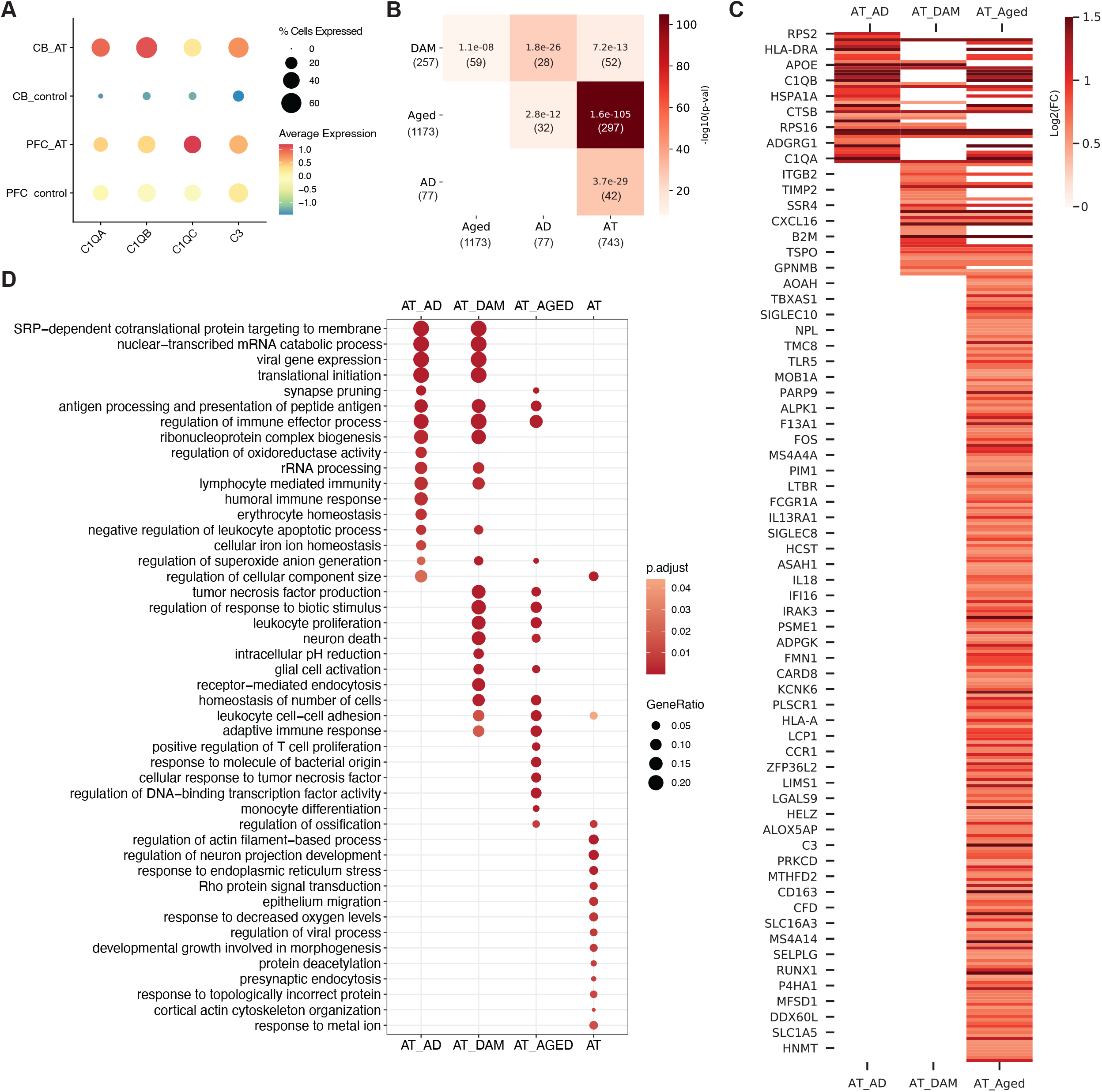
A-T cerebellar microglia share transcriptomic signatures with aging and neurodegenerative microglia. **A**. Average scaled expression of complement components *C1QA, C1QB, C1QC, C3* in microglia from A-T and control cerebellum (CB) and prefrontal cortex (PFC). **B**. Overlap between A-T microglia upregulated genes and human Alzheimer’s disease microglia markers (AD, Mathys et al., 2019), Disease Associated Microglia markers (DAM, Keren-Shaul et al., 2017), and human aging microglia markers (Aged, Olah et al., 2018). Overlap p-values from Fisher’s exact test shown in each cell. Color represents -log10(overlap p-value). Number of genes in each set and intersection shown in parentheses. **C**. Heatmap of A-T microglia log2 fold-changes (FDR<0.05) for overlapping microglia markers. AT_AD: A-T and Alzheimer’s disease microglia overlapping genes. AT_DAM: A-T and DAM overlapping genes. AT_AGED: A-T and human aging microglia overlapping genes. **D**. GO biological process enrichment of overlapping microglia markers. AT: A-T microglia only upregulated genes. Pathways with FDR<0.05 shown.

### Differential activation of microglia in A-T cerebellum versus PFC

To systematically compare gene expression changes in A-T cerebellum with A-T PFC, we identified genes and pathways that are more significantly changed in A-T cerebellum than A-T PFC in cell types common to the two brain regions (microglia, OPC, oligodendrocytes, astrocytes, endothelial cells) (Fig. S8, Tables S11-16). While both cerebellar and PFC microglia showed upregulation of genes involved in immune response in A-T, microglia were more strongly activated in the cerebellum compared to the PFC. Cerebellar A-T microglia had significantly higher upregulation of genes related to inflammatory processes (adaptive immune response, leukocyte proliferation, glial cell activation, tumor necrosis factor superfamily cytokine production) and neuronal death than A-T PFC microglia (Fig. 5A, Fig. S7).

**Figure 5.**
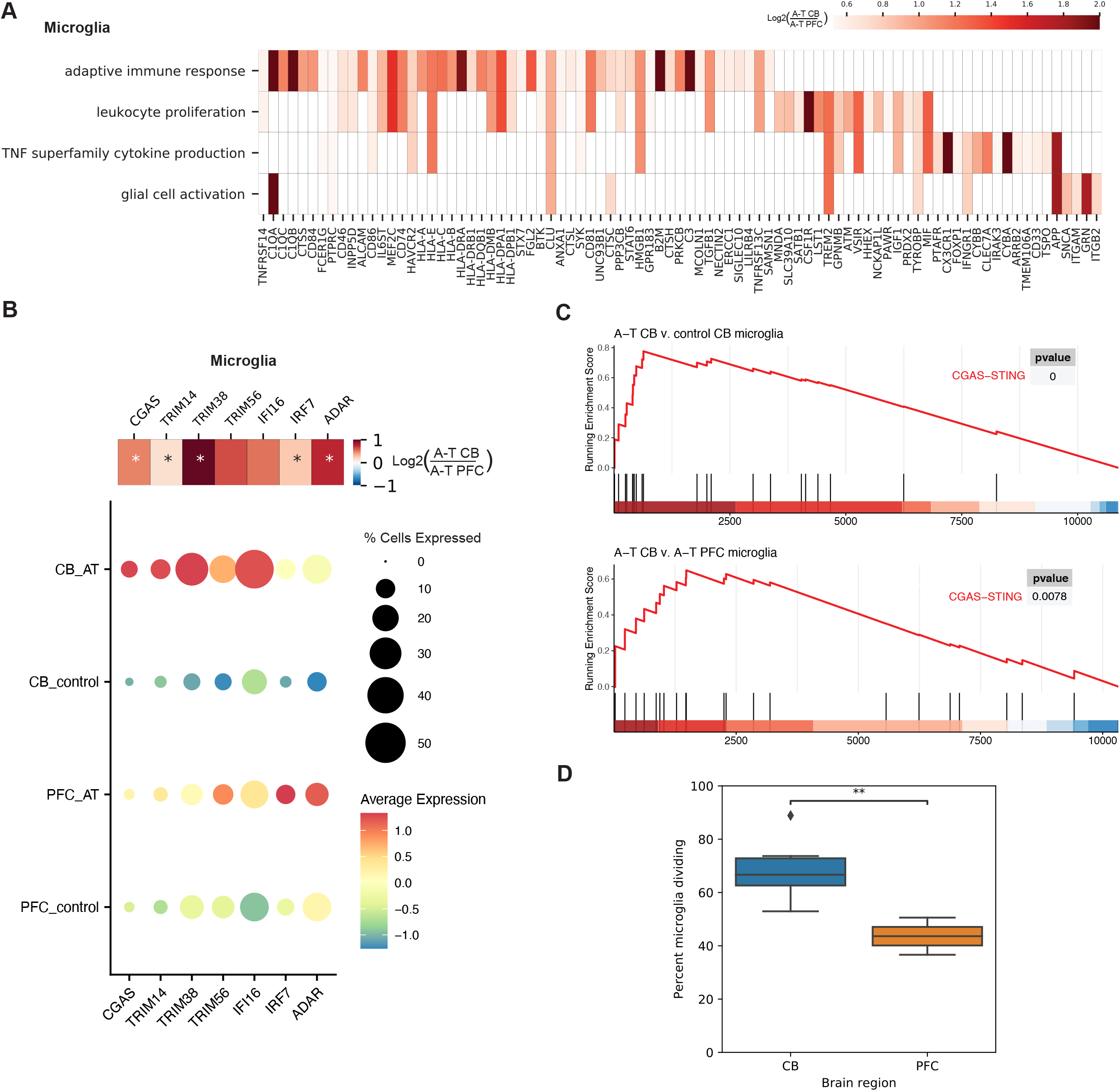
Stronger activation of microglia in A-T cerebellum compared to PFC. **A**. Heatmap showing enriched GO biological processes and log2 fold-changes of significant DEGs in each pathway with greater upregulation in A-T CB microglia than A-T PFC microglia. **B**. Dotplot of average scaled expression of CGAS-STING pathway genes in microglia of the cerebellum (CB) and PFC in A-T and control. Heatmap shows log2 fold-change of CGAS-STING pathway genes in A-T cerebellar microglia versus AT PFC microglia. *FDR<0.05. **C.** Gene Set Enrichment Analysis (GSEA) plot for the CGAS-STING pathway in A-T CB microglia versus control microglia (top), and A-T CB versus A-T PFC microglia (bottom). **D**. Percentage of microglia putatively in replicating phase in control cerebellum and PFC. **p-value <0.01, t-test.

Microglia can be activated by neuronal death, a mechanism common to multiple neurodegenerative conditions^44^. However, since PFC neurons do not degenerate in A-T, we hypothesized that A-T microglia in the PFC and cerebellum are cell-intrinsically activated. One potential mechanism of intrinsic activation is the accumulation of DNA damage due to *ATM* deficiency. Deficits in DNA repair caused by *ATM* deficiency are associated with increased cytosolic DNA^45^. Cytosolic DNA is recognized by the CGAS-STING pathway, which triggers an innate immune response by inducing type I interferons and inflammatory cytokines^46^. We found that the CGAS-STING pathway, including *CGAS* and activators of CGAS signaling—*TRIM14, TRIM38,* and *TRIM56*^47–49—were^ upregulated specifically in A-T microglia from the cerebellum and PFC, suggesting the presence of cytosolic DNA (Fig. 5B-C). Moreover, the CGAS-STING pathway showed significant enrichment among genes upregulated in A-T cerebellum versus A-T PFC, suggesting that CGAS-STING activation was more prominent in cerebellar microglia (Fig. 5C). CGAS-STING pathway genes (*TRIM14, TRIM38, IFI16, IRF7*) were also upregulated in aged human microglia but not AD or DAM gene sets, implicating DNA damage and cytoplasmic DNA as a common underlying cause of microglial activation in A-T and aging microglia, while AD microglia may be activated by another mechanism such as extrinsic stimuli (Table S17). Taken together, activation of the CGAS-STING pathway suggests that cytosolic DNA contributes to microglial activation that is more pronounced in AT cerebellum than PFC.

Self-DNA enters the cytosol in a mitosis dependent manner, through the formation and rupture of DNA-damage induced micronuclei^50,51^. Thus, we hypothesized that underlying differences in proliferation may explain the differential activation of CGAS-STING in cerebellar microglia compared to PFC microglia. We tested whether microglia in the cerebellum and PFC differ in proliferation rate by inferring the cell cycle status of each cell by the expression of canonical markers of G1, S, and G2 phases. We found that cerebellar microglia have an increased percentage of cells in replicating phases compared to cortical microglia (p-value < 0.01, t-test) (Fig. 5D). This is consistent with previous reports that cerebellar microglia have a higher turnover rate than cortical microglia, which may make them more vulnerable to accumulation of DNA damage due to *ATM* deficiency^52,53^. The differences in proliferation may underlie our observation that cerebellar microglia have a stronger CGAS-STING mediated immune response in A-T than cortical microglia.

### A-T patient iPSC-derived microglia demonstrate cell-intrinsic activation of NF-kB and type I interferon signaling

To understand whether microglia might contribute to neurodegeneration in A-T, we sought to determine when cerebellar microglia become activated in disease progression relative to Purkinje and granule cell death. We employed pseudotime analysis to understand the cell-type-specific cascades of molecular events across disease progression in A-T^57^. For each cell type, cells were ordered along a continuous trajectory that progressed from the control to diseased state (Fig. S8A). We performed differential analysis to identify genes that change as a function of pseudotime and clustered these genes by their pseudotime expression patterns. We then applied GO analysis to determine the enriched biological functions of each gene cluster. For microglia, the beginning of pseudotime (healthy state) had high expression of genes involved in cellular respiration, oxidative phosphorylation, and synaptic vesicle exocytosis, consistent with a high metabolism gene expression profile characteristic of cerebellar microglia^58^ (Fig. S8B). Over disease progression, upregulation of CGAS-STING pathway genes (i.e., *CGAS, TRIM14, TRIM38, TRIM56*) was followed by leukocyte activation, cytokine production, and innate immune response (Fig. S8B-C). In Purkinje neurons, disease progression was characterized first by a decline in genes related to calcium ion homeostasis (i.e., *ITPR1),* followed by late of a cluster of genes related to immune response and response to cytokine (i.e., *HSPA1A, B2M, ISG15)*, and apoptotic signaling pathway (i.e., *XBP1, BCL2, CASP2)* (Fig. S8D). Granule neurons first demonstrated a decrease in chromatin organization (i.e., *HDAC4, HDAC2, CHD5)* and glutamatergic synaptic signaling genes (i.e., *NRXN1, RELN, SYT1)*, followed by upregulation genes related to oxidative phosphorylation (i.e., *PINK1, NDUFS5, UQCRH)*, then upregulation of genes related to response to cytokine (i.e., *SP100, STAT6, ISG15*), response to unfolded protein (i.e., *HSPA1A, HSPE1, DNAJB1*), and programmed cell death (i.e., *IFI6, BCL2, XBP1*) at the end of pseudotime, suggesting that synaptic and metabolic dysfunction precedes granule neuron death in A-T (Fig. S8E).

We then compared molecular events between microglia, Purkinje, and granule cells along disease progression by aligning their pseudotime trajectories using dynamic time warping^59^. This revealed that the upregulation of inflammatory response and cytokine production genes in microglia (i.e., *C1QA, IL18, IFI16, CGAS, LTBR, TNFSF10)* occurs earlier in disease progression pseudotime than the onset of response to cytokine and apoptosis genes (*XBP1, MUC1, TFAP2A)* in Purkinje and granule cells (Fig. 6A-B). These findings suggest that microglial activation and inflammation precedes Purkinje and granule cell death in A-T and support a model of cell-intrinsic microglial activation in A-T.

**Figure 6.**
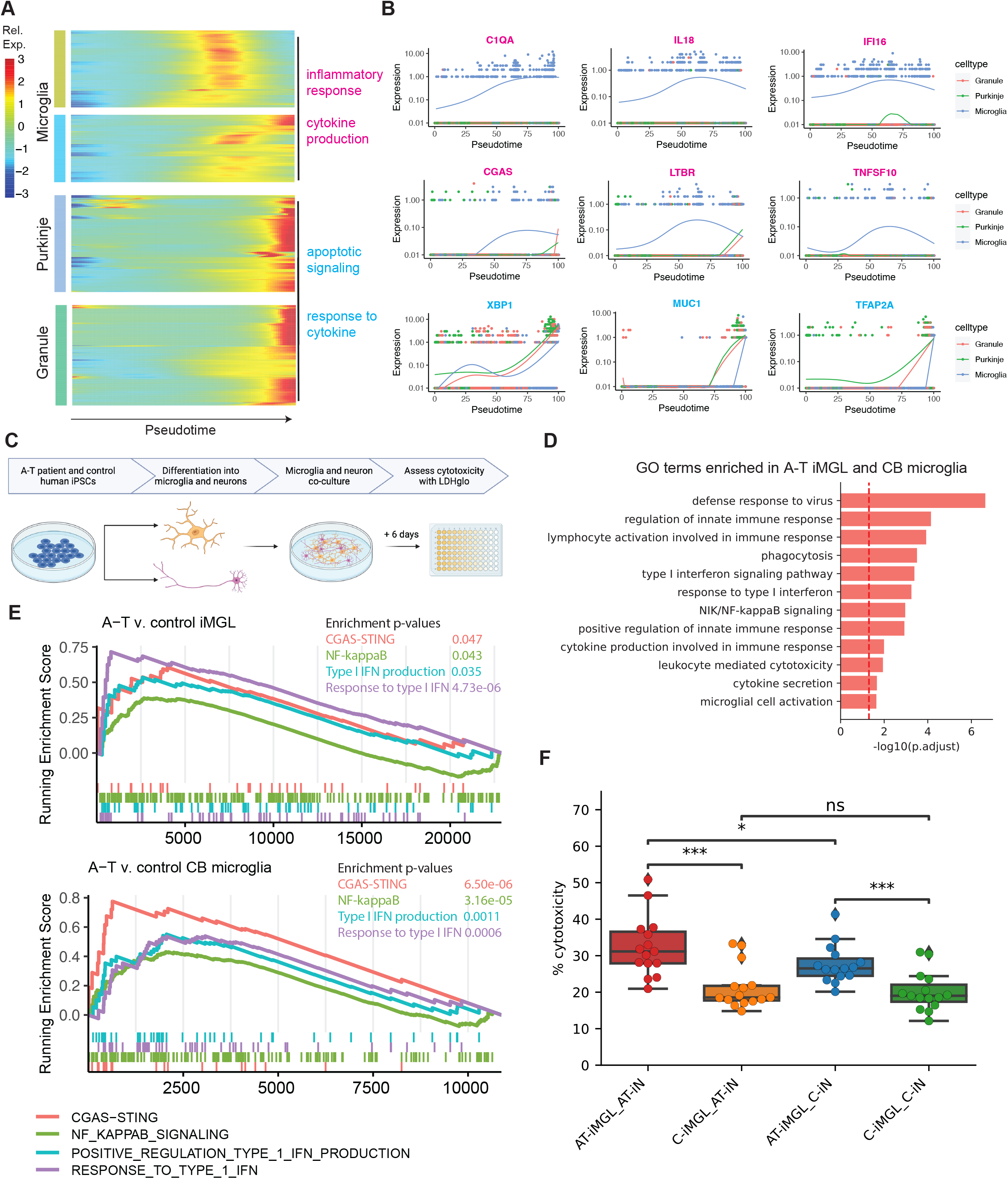
A-T patient iPSC-derived microglia reveal cell-intrinsic activation and increased cytotoxicity in neuronal co-cultures. **A**. Heatmap of genes that significantly change over alignedpseudotime in microglia, Purkinje, and granule neurons clustered by pseudotemporal expression patterns. Each cluster is annotated with enriched GO terms (FDR<0.05). **B**. Expression of select inflammatory genes (magenta) and apoptotic signaling and response to cytokine genes (cyan) over aligned pseudotime in microglia, Purkinje and granule neurons. **C**. Schematic of A-T and control iPSC-microglia (iMGL) and iPSC-neuron (iN) co-culture experiment. **D.** Gene Ontology (GO) terms enriched among genes upregulated in A-T iMGL and A-T cerebellar microglia versus control. **E.** Gene Set Enrichment Analysis (GSEA) plots for CGAS-STING, NF-kappaB, type I IFN production, and response to type I IFN pathways in A-T versus control iMGL (top) and A-T versus control CB microglia (bottom). **F.** Percent cytotoxicity of iMGL and iN co-cultures based on LDH-based cytotoxicity assay (bottom). C, PGP1 control. *p<0.05, ***p<0.001, ns, not significant, Mann-Whitney-U test.

To experimentally characterize microglial activation in A-T, we developed iPSC-based experimental models using A-T patients and controls (Fig. 1B, Fig. S9A). We generated human iPSCs from fibroblasts of an individual with A-T with compound heterozygous splice-altering variants in *ATM* (hg38 chr11:g.108332838C>T; chr11:g.108347266_108353792del27) and used the well-characterized PGP1 iPSC line as a control. We then differentiated microglia-like cells (iMGL) from the A-T patient and control iPSCs through a hematopoietic progenitor intermediate^54^ (Fig. S9A). The resulting iMGLs were a highly homogeneous population with >85% of cells expressing microglia markers CD45 and CD11B assessed by flow cytometry (Fig. S9B, Fig. S10A). Profiling of the iMGLs by RNA-sequencing showed that they expressed canonical markers of microglia (*AIF1, CX3CR1, CSF1R, SPI1, TREM2)* but did not express myeloid lineage (*MPO, KLF2),* neuronal (*MAP2, SOX2)*, and pluripotency (*POU5F1)* marker genes (Fig. S9C). Principal component analysis with transcriptomic profiles from A-T and control iMGLs along with our snRNA-seq data from human adult cerebellum and published snRNA-seq data from human fetal cerebellum^27^ confirmed that iMGLs cluster more closely with microglia from adult and fetal human cerebellum than with other brain cell types (Fig. S9D, Fig. S10B). Our clustering analysis also confirmed that iMGLs were most correlated with microglia in adult and fetal human cerebellum (Fig. S10C).

We performed differential gene expression analysis of A-T versus control iMGL, revealing upregulation of genes involved in cytokine secretion, phagocytosis, NF-kB signaling, and type I interferon signaling (Fig. S9E, Table S18-20). To compare A-T iMGL and A-T cerebellar microglia, we obtained genes upregulated in both and identified enrichment of immune related GO terms, including microglial cell activation, cytokine secretion, NF-kB, and type I interferon signaling pathway (Fig. 6D, Table S21). These results confirmed that A-T microglia grown in isolation take on an activated phenotype, demonstrating that this is a cell intrinsic property of microglia in A-T.

We hypothesized that cell-intrinsic activation of NF-kB and type I interferon signaling in A-T microglia could be due to cytoplasmic DNA triggering CGAS-STING, since a major consequence of CGAS-STING activation is initiation of NF-kB signaling and production of type I interferon related cytokines^55,56^. In support of cytoplasmic DNA as a potential trigger for microglia activation in A-T, A-T iMGL also showed positive enrichment of a CGAS-STING gene set (Fig. 6E, Table S22). The similar enrichment of CGAS-STING, NF-kB, and type I interferon pathways among genes upregulated in A-T iMGL and A-T postmortem microglia supports that A-T microglia are activated by a cell-intrinsic mechanism that may involve cytoplasmic DNA and CGAS-STING (Fig. 6E, Table S22).

### In-silico and in-vitro models of microglia-neuron interactions reveal that A-T microglia increase cytotoxicity

A-T Purkinje and granule neurons showed enrichment of response to cytokine genes late in disease progression, raising the possibility that cytokines produced by microglia induce molecular changes in Purkinje and granule neurons in A-T (Fig. 6A). Consistently, ligand-receptor analysis revealed increased cytokine signaling (*IL1B, IL7, IL4, IL15*), type I interferon signaling (*IFNA2),* CD40 signaling *(CD40LG),* and complement signaling *(C3, C4)* from microglia to granule and Purkinje neurons in A-T versus control (Fig. S9F, Fig. S11). Notably, several of these pro-inflammatory cytokines, including *IL1B* and *IFNA2,* have been shown to induce neurotoxicity^60–62^. This suggests that activated microglia in A-T have increased secretion of pro-inflammatory cytokines that may have neurotoxic consequences.

To test this hypothesis experimentally, we differentiated A-T patient and control iPSCs into neurons using overexpression of *Ngn2*^63^ and co-cultured them with A-T patient and control iMGLs (Fig. 6C). iPSC-derived neurons (iNs) transcriptionally correlated with human midgestational cerebellum, making them a relevant model for studying microglia-neuron interactions in A-T (Fig. S9D). We co-cultured iMGLs and iNs for six days and assessed cytotoxicity by lactate dehydrogenase (LDH) release. We found that co-culturing control iNs with A-T iMGLs was sufficient for increased cytotoxicity compared to co-culturing with control iMGLs (Fig. 6C, p-value<0.001). Moreover, A-T iMGLs co-cultured with A-T iNs demonstrated increased cytotoxicity compared to control iMGLs co-cultured with A-T iNs (Fig. 6F, p-value<0.001). A-T iMGLs co-cultured with A-T iNs had the highest levels of cytotoxicity, suggesting that A-T iNs are more vulnerable to A-T iMGLs. Taken together, these results indicate that A-T microglia have increased pro-inflammatory interactions with neurons and lead to increased cytotoxicity.

## DISCUSSION

In this study, we present an atlas of human cerebellar transcriptomes in health and degeneration at single-cell resolution. We identified major neuronal and glial cell types in the cerebellum and recovered known and novel transcriptomic signatures in A-T. We resolved cell-type-specific transcriptional perturbations in A-T, including the activation of pro-inflammatory pathways in microglia that was more pronounced in the cerebellum versus PFC. The activation of microglia in A-T PFC despite lack of cortical atrophy or neuronal loss suggests that A-T microglia are activated by a cell-intrinsic mechanism rather than in response to neurodegeneration, with brain region specific factors leading to stronger activation in the cerebellum. We found that genes in the CGAS-STING pathway, including *CGAS* itself, were significantly upregulated specifically in A-T microglia, with preferential activation in the cerebellum. Moreover, A-T patient iPSC-derived microglia cultured in the absence of neurons also demonstrated upregulation of genes involved in CGAS-STING, NF-kB, and type I interferon signaling, further suggesting a cell-intrinsic mechanism of activation. The CGAS-STING pathway is activated in response to cytosolic DNA, suggesting the presence of cytosolic DNA in A-T microglia. These results are consistent with recent reports of cytosolic DNA and CGAS-STING activation in microglia from rodent models of A-T^64,65^. Potential sources of cytosolic DNA include release of nuclear self-DNA^45,46^, mitochondrial DNA released by defective mitochondria^14,66,67^, reverse-transcribed transposable elements as observed in aging-related disorders^68^, and phagocytosis of nucleic acids from cellular debris. It will be of great interest for future work to identify the source of cytosolic DNA in A-T microglia, as it is a novel potential therapeutic avenue for modulating inflammation in A-T. Reducing inflammation in rodent models of A-T with ibuprofen led to decreased microglial activation and neuronal apoptosis, further highlighting microglia as a therapeutic target in A-T^69^.

Activated microglia have been intensely studied in the context of neurodegenerative disorders including Alzheimer’s disease, Parkinson’s disease, and ALS^70^. Our study highlights the involvement of microglia in human cerebellar degeneration as well. Through pseudotime analyses, we found that the rise in pro-inflammatory genes in A-T microglia precedes the onset of cytokine response, and apoptosis related genes in A-T Purkinje and granule neurons, and ligand-receptor analysis indicated increased pro-inflammatory signaling from microglia to neurons in A-T. These findings support a model of pathogenesis whereby cell-intrinsically activated microglia release neurotoxic cytokines and factors that trigger Purkinje and granule cell death. Recent work in human microglial cell lines demonstrated that persistent DNA damage due to loss of ATM leads to microglial activation and excessive phagocytosis of neurites, providing further validation for a cell-intrinsic mechanism of microglia activation that contributes to neuronal damage^10^. Rodent models of A-T also show evidence of neuroinflammation and microglial activation that leads to secretion of neurotoxic cytokines^64,65,71^ but why these models do not fully recapitulate cerebellar degeneration remains unknown. Potential reasons may be that the rodent cerebellum has an increased numerical density (number/mm^3^) of Purkinje cells compared to human cerebellum^72^, or Purkinje cell transcriptomic differences between rodent and human may underlie the different phenotypes^73,74^. Here, our data from human A-T patients and patient-derived iPSC models provides critical support that microglial activation, previously demonstrated *in vitro* and in rodent models, is also found in human A-T patient brains and may contribute to A-T disease mechanisms. The pathological nature of pro-inflammatory signaling from activated microglia is also supported by clinical observations that treatment with glucocorticoids reduced ataxia symptoms in A-T ^75–78^. The presence of activated microglia in A-T PFC without cortical neuron loss suggests that cerebellar Purkinje and granule neurons are more vulnerable to activated microglia. Moreover, the increased cytotoxicity of A-T iNs co-cultured with A-T iMGLs suggest that A-T iNs have increased vulnerability to microglia. It is highly possible that cell autonomous and non-cell autonomous effects of *ATM* loss in cerebellar neurons and microglia interact to cause A-T pathology. Given that microglia and other brain-resident macrophages share several transcriptomic markers, it is possible that the microglia cluster may include a smaller percentage of other brain-resident macrophages. As markers of subtypes of brain-resident macrophages become elucidated, future studies will be able to better distinguish these cell types in the brain. Moreover, it is possible that some of the transcriptional alterations identified in certain cell types reflect processes other than degeneration, such as compensatory changes in surviving cells. It will be important for future work to further dissect the developmental onset, contribution of cell type interactions, and cellular responses to cerebellar degeneration in A-T.

This study identifies cell-type-specific changes in A-T across multiple brain regions and serves as a resource as an adult human cerebellum vermis dataset in health and degeneration.

While granule and Purkinje neurons have been the most studied cell types in A-T due to their selective degeneration, this study highlights the central role of microglia in A-T cerebellar degenerative pathogenesis.

## Acknowledgements

We acknowledge the NIH NeuroBioBank for providing human tissues used in this study. We would like to thank Brad Margus from the AT Children’s Project for helpful scientific discussion. We also acknowledge Allison R. McLean and Jeffery R. Gulcher from Genuity Sciences for their assistance with snRNA-seq data coordination, Ronald Mathieu from HSCI-BCH Flow Cytometry Research Facility for his assistance with flow cytometry, and HSCI iPS Core Facility for their assistance with iPSC derivation. Illustrations were created with BioRender.com.

This work was supported by the National Institute of Health grants: K01 AG051791 (E.A.L), DP2 AG072437 (E.A.L), R00 AG054748 (M.A.L), AT Children’s Project (E.A.L, M.A.L.), Suh Kyungbae Foundation (E.A.L), Paul G. Allen Family Foundation (E.A.L), The American Federation for Aging Research (M.A.L. and A.M.J.), and Charles H. Hood Child Health Foundation (E.A.L., M.A.L). J.L. was supported by award T32GM007753 from the National Institute of General Medical Sciences. The content is solely the responsibility of the authors and does not necessarily represent the official views of the National Institute of General Medical Sciences or the National Institutes of Health.

## Author contributions

E.A.L., M.A.L., J.K., and J.L. conceptualized the study. M.A.L., A.J., T.W.C., and P.G.B. performed single-nucleus RNA-sequencing experiments. J.L and J.K analyzed the single-nucleus RNA-sequencing data. J.L and D.C performed iPSC-derived experiments. A.T. performed the immunohistochemistry analyses. J.L and J.P performed WGS analysis. E.AL., T.W.Y, and M.A.L. supervised the study. J.L. wrote the manuscript. All authors reviewed and revised the manuscript.

## Methods

### Isolation of single nuclei from fresh frozen tissue samples

All human postmortem brain tissue samples were obtained from the NIH NeuroBioBank. Samples were matched as closely as possible for age, sex, RIN, and PMI. All available neuropathology reports and medical records for each sample were obtained. Detailed sample information is in Table S1. All cerebellar vermis samples are from the same anatomical section (section 1) and include the entire vermis, and all cortex samples are from BA9/BA46, which are adjacent Brodmann areas. Sectioning was performed as described by the University of Maryland Brain and Tissue Bank Sectioning Protocol: “LEFT HEMISPHERE: CEREBRUM A cut is made just posterior to the cerebral peduncle and the midbrain/pons/cerebellum are removed as a unit from the left hemisphere (Cut #3). The remaining cerebrum is sectioned coronally, at approximate 1 cm intervals beginning from the frontal pole apex and proceeding caudally. As each section is isolated, it is gently rinsed with water, blotted dry, assigned a sequential numeric identifier (odd numbers only!), and placed in the freezing bath. The handling of sections is best aided by the use of a plastic spatula. Each frozen section is placed into individual plastic bags appropriately labeled and sealed. All bags are then stored in a −80 degree Centigrade freezer prior to shipping. Frozen sections of the cerebrum are identified as sections 1,3,5,7,9.

LEFT HEMISPHERE: CEREBELLUM The remaining cerebellum is placed in a vertical plane (its normal anatomic position) and sectioned at 0.5 to 0.6 cm intervals beginning from the medial surface (vermis) and moving laterally. Each resulting section is assigned a sequential identifier (odd numbers only!). Frozen sections of the cerebellum are identified as sections 1,3,5,7,9” (https://www.medschool.umaryland.edu/btbankold/Brain-Protocol-Methods/Brain-Sectioning---Protocol-Method-2/).

The nuclei isolation protocol was adapted from two previous publications^79,80^. Specifically, the protocol was performed as follows: all procedures were performed on ice or at 4°C. Approximately 100 mg of fresh frozen samples were processed using a Dounce homogenizer in 5 ml of tissue lysis buffer (0.32M sucrose, 5mM CaCl2, 3mM MgAc2, 0.1mM EDTA, 10mM Tris-HCl (pH 8), 0.1% Triton X-100, 1mM DTT). The homogenized solution was loaded on top of a sucrose cushion (1.8M sucrose, 3mM MgAc2, 10mM Tris-HCl (pH 8), 1mM DTT) and spun in an ultracentrifuge in an SW28 rotor (13,300 RPM, 2hrs, 4°c) to separate nuclei. After spinning, supernatant was removed and nuclei were resuspended (1% BSA in PBS plus 25ul 40U/ul RNAse inhibitor) then filtered through a 40um cell strainer. After filtration, nuclei were counted and diluted to a concentration of 1000 cells/ul.

### Droplet-based snRNA-seq

Droplet-based libraries were generated using the Chromium Single Cell 3’ v3 reagent kits (10x Genomics) according to manufacturer’s instructions. Resulting libraries were sequenced on an Illumina Novaseq at 150 paired-end reads with depth 20,000 reads/nucleus.

### snRNA-seq analysis

snRNA-seq data for each individual was preprocessed and aligned to the GRCh38-3.0.0 reference genome using CellRanger count. After alignment, ambient/background RNA was removed from the count matrix using CellBender remove-background (v0.2.0)^20^. The resulting count matrices were used to create Seurat objects per individual using Seurat v4^18^. Further quality control removed cells with less than 200 genes per cell, greater than 5000 genes per cell, and greater than 30% mitochondrial content. DoubletFinder^19^ was used to detect and remove doublets with an expected multiplet rate of 7.6% for ∼16,000 cells loaded per sample based on the 10x user guide.

SCTransform was used to perform normalization of expression values per sample^81^. After normalization, samples were integrated to align common clusters across individual datasets using Seurat’s integration method^18^. Dimensionality reduction was performed using RunPCA and RunUMAP. A shared nearest neighbor graph was constructed using FindNeighbors with dims 1:30, k.param 100, and cosine metric. Clusters were then identified using the Leiden algorithm at a resolution of 1.5.

Marker genes for each cluster were obtained using FindAllMarkers with a minimum log2 fold-change of 0.25. Cluster markers were used to assign cerebellar cell type identities based on known literature cell type markers. For annotating clusters in the prefrontal cortex, SingleR was used for reference-based annotation since reference datasets from the human prefrontal cortex are available^23^. Velmeshev et al., 2019 was used as the prefrontal cortex reference^13^.

To identify changes in cell type proportions in AT compared to controls, we used a Bayesian statistics approach to compositional data analysis previously described in the context of microbial ecology^24^. Compositional data has a negative covariance structure that is accounted for by the multinomial and Dirichlet probability distributions. We implemented Dirichlet multinomial modeling (DMM) using Hamiltonian Monte Carlo (HMC) using the R packages rstan and bayestestR (Stan Development Team (2020). “RStan: the R interface to Stan.” R package version 2.21.2, http://mc-stan.org/.)^82^. The input to the model was a matrix of counts where the rows correspond to replicates (individual samples) and the columns correspond to cell types. We then modeled the replicates using the multinomial distribution for the probability of each cell type per individual. The Dirichlet distribution was used to model the multinomial parameters. The prior on the Dirichlet parameters was another Dirichlet distribution with a fixed parameter alpha 10^-7^, which gives uniform cell type proportions in expectation. The resulting posterior distributions were Dirichlet distributions of the cell type proportions in AT and control. We then subtracted the posterior probability distribution (89% credible interval) of control from AT to see whether there are significant differences in relative cell type composition. A cell type shift in log2 abundance ratio (AT/control) was considered significant if the 89% credible did not include 0.

### Differential gene expression and Gene Ontology analysis

Differential gene expression analysis was performed on each cell type using edgeR^83^. The integrated Seurat object was subset to obtain Seurat objects for each cell type. The read counts were modeled and normalized using a negative binomial distribution with the trimmed mean of M-values (TMM) normalization method. To ensure robust signals, only genes that were expressed in at least 5% of one cell type were included in the analysis. The design matrix formula was ∼ disease.status + sex + age + cellular detection rate. Differentially expressed genes between AT and control were identified using the likelihood ratio test (glmLRT). Genes with FDR < 0.05 and a log2 fold-change greater than 0.50 were used for downstream analysis.

Pseudobulk differential gene expression analysis was performed using DESeq2^84^. To generate pseudobulk data, we took the sum of raw counts for each gene over all cells in each sample, resulting in a gene by sample counts matrix. The design matrix was specified as ∼ sex + age + disease.status. Differentially expressed genes between AT and controls were identified after running lfcShrink. Genes with FDR < 0.05 were considered statistically significant.

Gene Ontology (GO) enrichment analysis was performed on the upregulated and downregulated differentially expressed genes in each cell type using clusterProfiler^85^. GO biological processes with FDR < 0.05 were considered significant.

Gene Set Enrichment Analysis was performed on differentially expressed genes ordered by log2 fold-change (descending order) with the function GSEA() with default settings from clusterProfiler. Human gene sets of interest were obtained from the Molecular Signatures Database (MSigDB)^86,87^ and are listed in Supplementary Table 21.

CellChat^88^ was used to perform ligand-receptor analysis in order to identify differential cell-cell communication in AT compared to control. Differentially active signaling pathways with FDR<0.05 were considered significant.

### Cell Cycle Scores

AddModuleScore calculates the average expression of a gene set in each cell, subtracted by the average expression of a control gene set, as previously described^89^. A positive score means the module of genes is expressed more highly in a particular cell than expected by chance. Cell cycle phase scores were calculated using this method based on canonical cell cycle markers using the Seurat function CellCycleScoring, which then outputs the predicted classification (G1, S, or G2M phase) for each cell.

### Disease gene analysis

Known monogenic cerebellar disease genes were obtained from the Online Mendelian Inheritance in Man (OMIM) database. The mouse cerebellum data was obtained from Saunders et al., 2018^90^. To determine the correlation of cerebellar disease gene expression between humans and mice, cosine similarity was calculated between the human and mouse expression vectors of the genes for each cell type.

### Pseudotime analysis

Pseudotime trajectory analysis was performed using Monocle2^57^. First, Seurat objects for each cell type (Microglia, Purkinje, and granule cells) were converted into the Monocle2 compatible cell dataset form using as.CellDataSet. Cells were clustered in an unsupervised manner using clusterCells. The top 1000 differentially expressed genes between AT and control in each cell type were used as the ordering genes to order cells along the pseudotime trajectory. The root state of the trajectory was defined as the cluster with the highest proportion of control cells. Genes that change as a function of pseudotime were identified using differentialGeneTest with the model formula specified as ∼sm.ns(Pseudotime), which fits a natural spline to model gene expression as a smooth nonlinear function of pseudotime. Genes that change over pseudotime with an FDR<0.05 were considered significant. These genes were then clustered according to their pseudotime expression profiles. GO enrichment analysis was performed on each cluster using clusterProfiler to identify the biological processes that co-vary across pseudotime.

To align the pseudotime trajectories between cell types, the method dynamic time warping (DTW) was applied as previously described^59^. DTW aligns similar temporal processes that differ in length by calculating the optimal matching between points in each sequence. For each cell type, ordered cells in pseudotime were binned into 100 “pseudocells” and the average expression of each gene was calculated within each pseudocell. The resulting gene x pseudocell matrices were the input for the dtw package in R. The Purkinje cell matrix was set as the reference and the other cell type was the query. The cost matrix was calculated as 1-corr(M1, M2), where corr(M1, M2) is the cosine similarity matrix between cell type 1 matrix (M1, reference) and cell type 2 matrix (M2, query). The query cell type pseudotime values were then warped into the reference to obtain aligned pseudotimes.

### Whole Genome Sequencing and Variant Calling

DNA was isolated using the Qiagen DNA Mini kit (Qiagen cat. no. 51304) according to the protocol for tissues. Approximately 25mg of fresh frozen tissue was minced into small pieces. Tissue was transferred to a 1.5ml microcentrifuge tube and 180µl of Buffer ATL as added. 20ul of proteinase K was added prior to 4 hours of agitation at 56 degrees centigrade on a thermomixer (1400 RPM). DNA isolation proceeded as written in the protocol with the inclusion of the optional RNase A step. Libraries were prepared with the TruSeq DNA PCR-free library kit (Illumina). Libraries were sequenced at 30X coverage with Illumina NovaSeq 6000 S4, 2 x 150 bp. WGS data for each individual was processed using GATK4^91^. Specifically, we used the DRAGEN-GATK whole genome germline pipeline for variant discovery with the WGS_Maximum_Quality mode on Terra (terra.bio). Reads were aligned using hg38 as the reference genome. Resulting VCF files were annotated using ANNOVAR^92^. Sample information and variants are listed in Table S5.

### Immunostaining and Imaging

Postmortem human formalin fixed vermis samples from four subjects diagnosed with Ataxia Telangiectasia and three age matched controls were obtained from the University of Maryland Brain and Tissue Bank (Table S9). The samples were embedded in paraffin by the UMCCTS Biospecimen, Tissue and Tumor Bank, and sectioned on a LEICA RM2125 microtome. Sections were cut at 20 µm thickness, placed on uncharged untreated Fisherfinest premium microscope slides (product number 12-544-7), and dried overnight at 37°C. Eight sections (four control and four with AT pathology) were selected at random, deparaffinized with xylene and rehydrated in a graded ethanol wash series. This was followed by antigen retrieval, done by heating citrate buffer (pH 6.2) to 90°C and leaving the slides to soak for 45 minutes, then allowing the solution and slides to cool to room temperature. The samples were then subjected to brief PBS washes before application of a primary antibody solution consisting of blocking solution, 10% triton in 10x PBS for permeabilization, and primary antibody. Primary antibodies used included CD11B/Integrin Alpha M Monoclonal Antibody (Proteintech, product number 66519-1-Ig, concentration 1:100), and Anti-GFAP antibody produced in rabbit (Millipore Sigma product number HPA056030, concentration 1:250). The samples were incubated at 4°C overnight. The following day the samples were washed for an hour and a half in cold 1x PBS before application of secondary antibody solution consisting of blocking solution, 10% triton, and Donkey anti-Mouse Cross-Adsorbed Secondary Antibody, Alexa Fluor plus 555 (ThermoFisher Scientific, Invitrogen, product number A32773, concentration 1:125) and Donkey anti-Rabbit Cross-Adsorbed Secondary Antibody Alexa Fluor plus 488 (ThermoFisher Scientific, Invitrogen, product number A32790, concentration 1:125). After incubating overnight in the dark at 4°C the samples were washed in 1x PBS for an hour and a half in the dark. The tissue was stained for DAPI (ThermoFisher Scientific, Product Number 62248, concentration 1:10,000 in 1x PBS), washed and then treated with TrueBlack lipofuscin autofluorescence quencher (Biotium, product number 23007) according to the manufacturer’s protocol. The samples were subjected to 30 minutes of washes in Barnstead water before being allowed to air dry. The tissue was mounted in Everbrite (Biotium, product number 23003) with Platinum Line cover glass (FisherScientific, Product Number 15-183-89) and left to cure overnight. Imaging was performed on a Zeiss LSM700 inverted microscope using ZEN Black 2012 Software.

### Maintenance and culture of iPSCs

Fibroblasts from an individual with A-T (2 years old, female) were reprogrammed into iPSCs using non-integrative Sendai virus. The patient had the following compound-heterozygous variants in *ATM:* hg38 chr11:g.108332838C>T; chr11:g.108347266_108353792del27. Control iPSCs from Personal Genome Project participant 1 (PGP1) were used as the control line. Both iPSC lines were mycoplasma negative, karyotypically normal, and expressed pluripotency markers *OCT4, SOX2, NANOG, SSEA4, TRA-1-60.* iPSCs were maintained in 6-well or 10 cm plates (Corning) coated with LDEV-free hESC-qualified Matrigel (Corning Catalog #354277) in feeder-free conditions with complete mTeSR-plus medium (STEMCELL Technologies) in a humidified incubator (5% CO2, 37°C). iPSCs were fed fresh media daily and passaged every 3-4 days.

### Differentiation of iPSCs to Hematopoietic Progenitor Cells

Human iPSC-derived hematopoietic progenitors (HPC) were generated using the STEMdiff Hematopoietic Kit (STEMCELL Technologies) according to the manufacturer instructions based on a protocol by Abud et al., 2017^54^. Briefly, iPSCs were dissociated into aggregates using Gentle Cell Dissociation Reagent (STEMCELL Technologies) and 40-80 aggregates were plated per well on Matrigel coated 12-well plates. Cells were maintained in Medium A to induce a mesoderm-like state until day 3. From day 3 to day 12, cells were maintained in Medium B with half-media changes every 2 days to promote further differentiation into hematopoietic progenitor cells. To confirm the differentiation from iPSC to HPC, we performed flow cytometry on day 12 at the end of HPC differentiation. Cells were harvested and incubated with CD45-APC (1 µl per 50,000 cells in 100 µL, STEMCELL Technologies Catalog #60018AZ) and CD34-PE (1 µl per 50,000 cells in 100 µL, STEMCELL Technologies Catalog #60119PE) antibodies for 15 minutes in the dark at 4C. In addition to double-stained samples, we included unstained samples, Fluorescence Minus One (FMO) controls, and compensation single-stain controls using OneComp eBeads (Thermo Scientific Catalog #01111142). Samples were washed twice with FACS buffer (Phosphate Buffered Saline (PBS) with 2% FBS) and transferred to 5 ml Round Bottom Polystyrene FACS tubes. Flow cytometry was performed at the HSCI-BCH Flow Cytometry Research Facility using the BD FACSAria II. Analysis was performed using FlowJo software.

### Differentiation of Hematopoietic Progenitor Cells to Microglia

Human iPSC-derived microglia (iMGL) were generated from human iPSC-derived hematopoietic progenitors using the STEMdiff Microglia Differentiation Kit and Microglia Maintenance Kit (STEMCELL Technologies) according to manufacturer instructions. Briefly, hematopoietic progenitor cells from day 12 of the STEMdiff Hematopoietic Kit protocol were collected and transferred to a Matrigel coated 6-well plate (200,000 cells per well) with STEMdiff Microglia Differentiation Medium. From day 0 to 24, 1 mL Microglia Differentiation Medium was added to each well every other day. After day 24, cells were maintained in STEMdiff Microglia Maturation Medium with half-medium volume added every other day. To confirm the differentiation from HPC to iMGL, we performed flow cytometry on day 24 at the end of microglia differentiation. Cells were harvested and blocked with CD32 antibody (2 µl per 50,000 cells in 100 µL, STEMCELL Technologies Catalog #60012) for 15 minutes at 4C. Samples were then incubated with CD45-APC (1 µl per 50,000 cells in 100 µL, STEMCELL Technologies Catalog #60018AZ) and CD11B-FITC (1 µl per 50,000 cells in 100 µL, STEMCELL Technologies Catalog #60040FI) antibodies for 15 minutes in the dark at 4C. In addition to double-stained samples, we included unstained samples, FMO controls, and compensation single-stain controls using OneComp eBeads (Thermo Scientific Catalog #01111142). Samples were washed twice with FACS buffer (Phosphate Buffered Saline (PBS) with 2% FBS) and transferred to 5 ml Round Bottom Polystyrene FACS tubes. Flow cytometry was performed at the HSCI-BCH Flow Cytometry Research Facility using the BD FACSAria II. Analysis was performed using FlowJo software.

### RNA-sequencing of A-T and control iMGL

RNA was harvested from day 29 iMGL samples using the PureLink RNA Mini kit (Life Technologies Catalog # 12183018A). Libraries were prepared with the Illumina Stranded mRNA prep. Sequencing was performed on Illumina HiSeq, 2×150bp configuration at approximately 20X depth per sample. Reads were aligned to hg38 using STAR v2.7.9a followed by processing with featureCounts to obtain a gene-level counts matrix. Differential gene expression analysis was performed using DESeq2 with control set as the reference condition and results coefficient set as ‘condition_AT_vs_control’.

### Differentiation of iPSCs to neurons

Human iPSC-derived neurons (iNs) were generated from iPSCs transduced with the lentiviruses Tet-O-Ngn2-Puro and FUW-M2rtTA based on a previously published protocol^63^. On day -1 of differentiation, iPSC colonies were dissociated into single-cells using Accutase (STEMCELL Technologies) and 4-8 million cells were plated on Matrigel coated 10 cm dishes with complete mTeSR plus medium supplemented with Y-27632 (10 uM). On day 0, cells were fed N2 media (DMEM/F-12 media, 1X N2, 1X Nonessential Amino Acids) supplemented with doxycycline (2 µg/mL), BDNF (10 ng/mL), NT3 (10 ng/mL), and laminin (0.2 µg/mL). On day 1, media was replaced with N2 media with puromycin (1 µg/mL) in addition to the supplements listed above. On day 2, cells were fed B27 media (Neurobasal media, 1X B27, 1X Glutamax) supplemented with puromycin (1 µg/mL), doxycycline (2 µg/mL), Ara-C (2 uM), BDNF (10 ng/mL), NT3 (10 ng/mL), and laminin (0.2 µg/mL). On day 3, cells were dissociated into single cells using Accutase and replated onto polyethylenimine/laminin coated 96-well plates (10,000 cells/well) with B27 media supplemented with Y-27632 (10 µM), doxycycline (2 µg/mL), Ara-C (2 µM), BDNF (10 ng/mL), NT3 (10 ng/mL), and laminin (0.2 µg/mL). On day 5, cells were fed with Conditioned Sudhof Neuronal Growth Medium (1:1 ratio of Astrocyte Conditioned Media and Neurobasal Media, 1X B27, Glutamax, NaHCO3, and transferrin) supplemented with BDNF (10 ng/mL), NT3 (10 ng/mL), and laminin (0.2 µg/mL).

### Co-culture of iMGL and iNs

Day 29 iMGL were added to Day 6 iNs plated in a 96-well plate at a 1:5 ratio (2,000 iMGL:10,000 neurons per well) with 15 independent wells per co-culture condition (A-T iMGL/A-T iN; A-T iMGL/control iN; control iMGL/A-T iN; control iMGL/control iN). The co-cultures were maintained for 6 days.

### Cytotoxicity assay

Cytotoxicity of co-cultures were assessed with the LDH-Glo Cytotoxicity Assay (Promega Catalog #J2380) according to the manufacturer instructions. 5 µl of medium from each co-culture well was diluted in 95 µl LDH Storage Buffer (200 mM Tris-HCl pH 7.3, 10% Glycerol, 1% BSA). LDH activity was measured by combining 10 µl LDH Detection Reagent with 10 µl diluted sample in a 384-well white opaque-walled assay plate (Greiner Bio-One Catalog # 784080) after incubating for 45 minutes at room temperature. LDH standards were prepared according to manufacturer instructions. Fresh culture medium was used as a no-cell control. To generate a maximum LDH release control, 2 µl of 10% Triton-X 100 was added per 100 µl to 96-wells containing iNs for 10 minutes followed by sample collection in LDH storage buffer as described above. Plate was read using the Spectramax iD5 plate reader with the luminescence read mode. Percent cytotoxicity was calculated according to the kit’s protocol as % Cytotoxicity = 100 × (Experimental LDH Release – Medium Background) / (Maximum LDH Release Control – Medium Background).

### Statistics and reproducibility

No statistical methods were used to predetermine sample sizes. All statistical tests were performed in R (v4.0.1) or Python (v3.8) with multiple hypothesis correction using the Benjamini-Hochberg procedure (FDR<0.05) unless otherwise specified. Statistical tests are noted in figure legends, and include Fisher’s exact test, Student’s t-test and Mann-Whitney U-test.

## Supplementary Figure Legends

**Fig. S1**. Pathogenic *ATM* variants in A-T cases. Alignment tracks of whole genome sequencing (WGS) reads at the *ATM* locus showing pathogenic variants in A-T cases.

**Fig. S2**. Quality control information for snRNA-seq data from cerebellum and PFC. **A.** Transcript (nCount_RNA), gene (nFeature_RNA), percent transcripts from mitochondrial genes, and percent transcripts from ribosomal genes per cerebellar sample before quality control filtering. **B.** Transcript (nCount_RNA), gene (nFeature_RNA), percent transcripts from mitochondrial genes, and percent transcripts from ribosomal genes per cerebellar sample after quality control filtering. **C.** UMAP plots of cerebellar data showing integration of samples, sex, age, and disease status. **D**. Stacked barplots showing proportion of each cell type per cerebellar sample. **E.** Posterior distribution of log2 ratio of cell type proportions in A-T cerebellum after 500 random permutations. **F**. Transcript (nCount_RNA), gene (nFeature_RNA), percent transcripts from mitochondrial genes, and percent transcripts from ribosomal genes per PFC sample before quality control filtering. **G.** Transcript (nCount_RNA), gene (nFeature_RNA), percent transcripts from mitochondrial genes, and percent transcripts from ribosomal genes per PFC sample after quality control filtering. **H**. UMAP plots of cerebellar data showing integration of samples, disease status, sex, and age. **I**. Stacked barplots showing proportion of each cell type per PFC sample. **J.** Posterior distribution of log2 ratio of cell type proportions in A-T PFC after 500 random permutations.

**Fig. S3**. snRNA-seq data of A-T PFC. **A.** UMAP plot of major cell types in AT and control human PFC, downsampled to 10,000 cells per condition only for visualization purpose. **B.** Relative abundance of cell types in AT versus control PFC, shown as the posterior distribution of Log2(proportion in AT/proportion in control) with 89% credible interval. Red bars highlight credible intervals that do not overlap 0. Bolded cell type labels indicate a significant difference in relative abundance. **C.** Differentially expressed genes (DEGs) in each cell type with FDR<0.05, |Log2FoldChange|>0.50. Each dot represents a significantly differentially expressed gene.

**Fig. S4.** Enriched expression of monogenic cerebellar disease genes in specific cell types. **A.** Dotplot of average scaled expression of *ATM* in cell types of the adult human cerebellum and prefrontal cortex, and **B**. developing human cerebellum (Aldinger et al., 2021). Microglia have the highest expression of *ATM* out of all cell types in the cerebellum and PFC. Size of dot represents percentage of single cells expressing the gene. Expression scaled to mean of 0 and standard deviation of 1. **C**. Heatmap of cerebellar ataxia disease gene expression across cell types in control human cerebellum. Color represents centered and scaled log-normalized expression.

*Enrichment p-value <0.05. OPC: oligodendrocyte precursor cell, IN-SV2C: SV2C expressing inhibitory neuron; IN-PV: PVALB expressing inhibitory neuron; IN-SST: SST expressing inhibitory neuron; L5/6-CC: layer 5/6 cortico-cortical neuron; L5/6: layer 5/6 neuron; L4: layer 4 neuron; L2/3: layer 2/3 neuron; 01-PC: Purkinje cells; 02-RL: Rhombic lip; 03-GCP: Granule cell progenitors; 04-GN: Granule neurons; 05-eCN/UBC: Excitatory cerebellar nuclei neurons/Unipolar brush cells; 06-iCN: Inhibitory cerebellar nuclei neurons; 07-PIP: PAX2+ interneuron progenitors; 08-BG: Bergmann glia; 09-Ast: Astrocytes; 10-Glia: Glia; 11-OPC: Oligodendrocyte precursor cells; 12-Committed OPC: Committed oligodendrocyte precursor cells; 13-Endothelial: Endothelial cells; 14-Microglia: Microglia; 15-Meninges: Meninges; 16-Pericytes: Pericytes; 17-Brainstem: Brainstem; 18-MLI: Molecular layer interneurons; 19-Ast/Ependymal: Astrocytes/ependymal cells; 20-Choroid: Choroid plexus; 21-BS Choroid/Ependymal: Choroid plexus/ependymal cell; SCA: spinocerebellar ataxia; SCAR: spinocerebellar ataxia, recessive; SCAN: spinocerebellar ataxia, autosomal recessive, with axonal neuropathy; CAMRQ: cerebellar ataxia, impaired intellectual development, and dysequilibrium syndrome.

**Fig. S5**. Gene Ontology enrichment of upregulated and downregulated genes across cell types in A-T cerebellum. **A.** Heatmap of enriched pathways among upregulated DEGs in cell types from A-T cerebellum. **B.** Heatmap of enriched pathways among downregulated DEGs in cell types from A-T cerebellum. Color represents z-score of pathway significance (-log10(p. adjusted)) and only significant pathways (p. adj <0.05) are colored.

**Fig. S6.** Dysregulation of hereditary ataxia genes in A-T. **A.** Heatmap showing log2 fold-change of hereditary ataxia genes in A-T cerebellum. *FDR<0.05. *ATM* expression increased in several cell types in A-T cerebellum, suggesting that lack of *ATM* function induces compensatory increases in transcription of the *ATM* locus. **B.** Enriched Gene Ontology (GO) pathways among disease genes with enriched expression in Purkinje cells and downregulated in A-T Purkinje cells. **C**. Overlap between *HDAC4* neuronal target genes and downregulated DEGs in A-T cerebellum.

**Fig. S7.** Enrichment of pathways among genes with greater dysregulation in A-T cerebellum than A-T PFC. **A**. Heatmap of enriched pathways among DEGs upregulated in AT CB v. AT PFC. **B.** Heatmap of enriched pathways among DEGs downregulated in AT CB v. AT PFC. Color represents z-score of pathway significance (-log10(p. adjusted)) and only significant pathways (p. adj <0.05) are colored.

**Fig. S8**. A-T patient iPSC-derived microglia reveal cell-intrinsic activation of NF-kappaB and type I interferon pathways. **A**. Schematic for generation of iPSC-derived microglia (iMGL) from human A-T patient and control iPSCs. PGP1, Personal Genome Project 1; HPC: hematopoietic progenitor cells. **B.** Flow cytometry analysis of CD45 and CD11B co-expression in control iMGLs 24 days post-differentiation. **C**. Expression of microglia (*AIF1, CX3CR1, CSF1R, SPI1, TREM2*), myeloid lineage (*MPO, KLF2*), neuronal (*MAP2, SOX2*), and iPSC (*POU5F1*) marker genes in A-T and control iMGLs. Each column represents data from iMGL differentiated in an independent well**. D**. Principal component analysis plot of A-T and control iMGL (iMGL-AT/control) and adult cerebellar cell type pseudobulk transcriptomic profiles derived from snRNA-seq in this study (AT/control-microglia) and fetal cerebellar cell type pseudobulk transcriptomic profiles derived from snRNA-seq data in Aldinger et al., 2021 (Fetal-microglia). **E**. Volcano plot for log2 fold change in gene expression in A-T versus control iMGL. Representative genes associated with cytokine secretion (purple), NIK/NF-kappaB (green), phagocytosis (red), and type I interferon (IFN) (blue) are highlighted. Horizontal dash line represents FDR=0.05. **F.** Ligand-receptor pairs with increased communication probability from microglia to granule or Purkinje neurons in A-T cerebellum versus control. Dot color represents communication probability (strength of signaling). Dot size represents significance of ligand-receptor pair and pairs with adjusted p-value <0.01 shown.

**Fig. S9**. Differentiation of A-T patient and control human iPSCs into microglia and neurons. **A.** Flow cytometry analysis of CD45 and CD11B co-expression in A-T and control iMGLs 24 days post-differentiation from 2 independent differentiation batches. **B**. Principal component analysis plot of A-T and control iMGL and adult cerebellar cell type pseudobulk transcriptomic profiles derived from snRNA-seq in this study and fetal cerebellar cell type pseudobulk transcriptomic profiles derived from snRNA-seq data in Aldinger et al., 2021. **C.** Heatmap of Spearman Correlation Coefficients between human adult and fetal cerebellar cell type transcriptome profiles and iMGL transcriptome profiles. **D**. Heatmap of Spearman Correlation Coefficients between human control (PGP1) iPSC-derived neurons at day 6 of differentiation transcriptomes and human developing cerebellum (CB) and cerebellar cortex (CBC) transcriptomes from BrainSpan: Atlas of the Developing Human Brain. iMGL-AT, A-T iPSC-derived microglia. iMGL-WT, control iPSC-derived microglia.

**Fig. S10**. Pseudotime analysis of disease progression reveals early microglia activation. **A**. Disease progression pseudotime trajectory of A-T cerebellar microglia, colored by disease status (red: A-T, blue: control), or pseudotime (healthy to diseased). **B**. Heatmap of genes that significantly change over pseudotime in microglia, clustered by pseudotemporal expression patterns. Each cluster is annotated with enriched GO terms (FDR<0.05). **C**. Expression of *CD11B, CGAS, TRIM14, TRIM38,* and *TRIM56* over pseudotime in microglia. **D**. Heatmap of genes that change over pseudotime in Purkinje neurons, clustered by pseudotemporal expression patterns. Each cluster is annotated with enriched GO terms (FDR<0.05). **E**. Heatmap of genes that change over pseudotime in granule neurons, clustered by pseudotemporal expression patterns. Each cluster is annotated with enriched GO terms (FDR<0.05). Heatmaps in B, D, E depict centered and scaled expression.

**Fig. S11**. Altered cell-cell communication in A-T cerebellum. **A.** Relative information flow for signaling pathways enriched in A-T or control cerebellum. Information flow for each pathway is calculated as the sum of the communication probability among all pairs of cell groups. **B.** Ligand-receptor pairs with increased communication probability between microglia, granule, and Purkinje neurons in A-T cerebellum compared to control. Dot color represents communication probability (strength of signaling) and dot size represents significance of ligand-receptor pair (adjusted p-value). Empty space represents a communication probability of zero.

## Supplementary Tables

Table S1: Detailed sample information

Table S2: Sample summary statistics

Table S3: Cell counts

Table S4: Neuropathology reports

Table S5: Pathogenic *ATM* variants identified by WGS

Table S6: OMIM genes list

Table S7: A-T CB DEGs per cell type

Table S8 GO terms enriched among A-T CB upregulated genes

Table S9: Cases used for immunohistochemistry

Table S10: GO terms enriched among A-T CB downregulated genes

Table S11: AT PFC DEGs per cell type

Table S12: GO terms enriched among A-T PFC upregulated genes

Table S13: GO terms enriched among A-T PFC downregulated genes

Table S14: AT CB versus PFC DEGs

Table S15 GO terms enriched among A-T CB versus PFC upregulated genes

Table S16: GO terms enriched among A-T CB versus PFC downregulated genes

Table S17: AD/Aged/DAM MICROGLIA gene sets

Table S18: iMGL DEGs

Table S19: GO terms enriched among A-T iMGL upregulated genes

Table S20: GO terms enriched among A-T iMGL downregulated genes

Table S21: GO terms enriched among upregulated genes shared by A-T iMGL and A-T CB

Table S22: GSEA statistics and CGAS, NFkB, type I IFN gene lists

