## Supplemental Figures for "ATM-deficiency induced microglial activation promotes neurodegeneration in Ataxia-Telangiectasia"

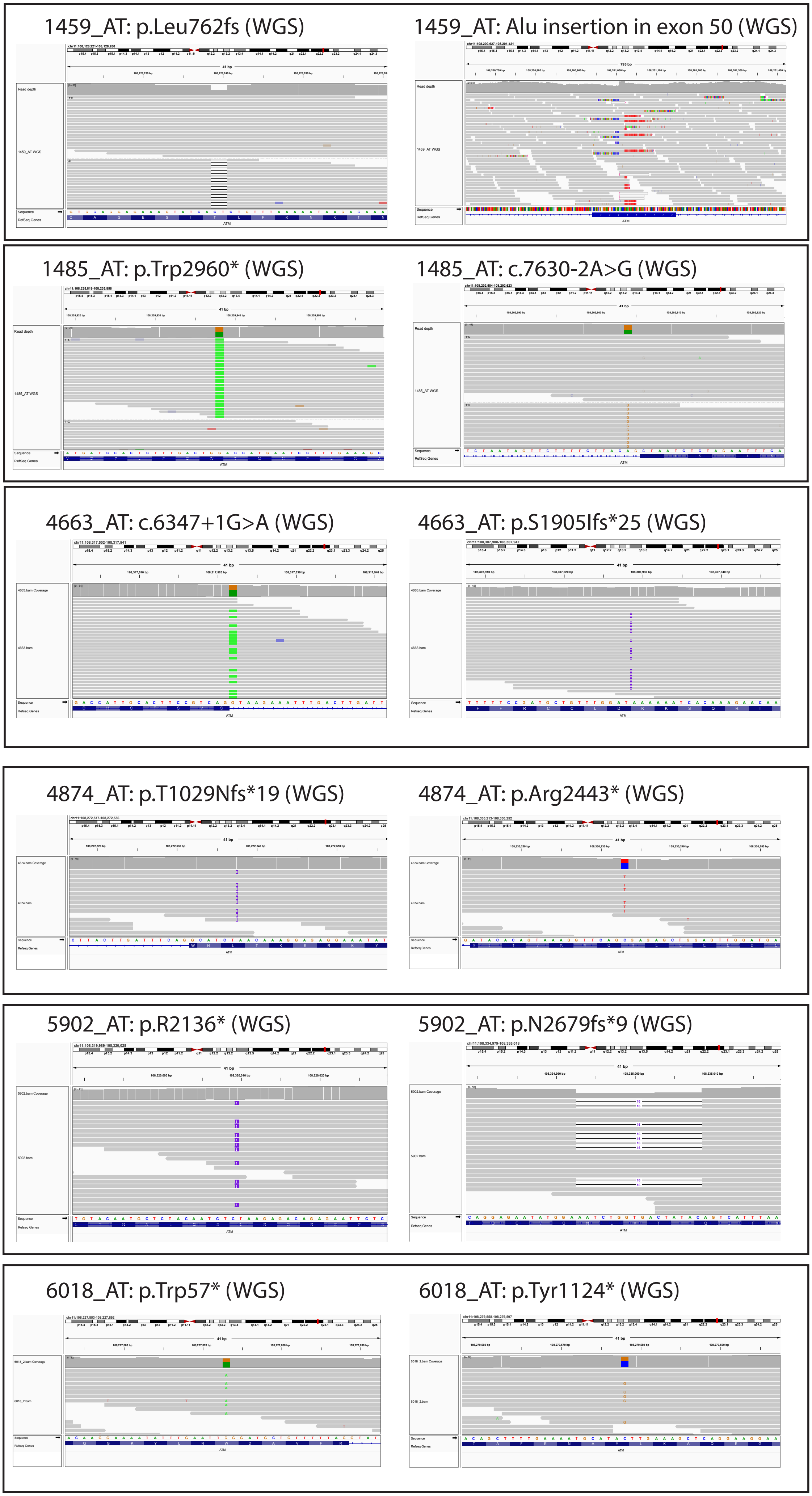
1459_AT: p.Leu762fs (WGS)
1459_AT: Alu insertion in exon 50 (WGS)
1485_AT: p.Trp2960* (WGS)
1485_AT: c.7630-2A>G (WGS)
4663_AT: c.6347+1G>A (WGS)
4663_AT: p.S1905Ifs*25 (WGS)
4874_AT: p.T1029Nfs*19 (WGS)
4874_AT: p.Arg2443* (WGS)
5902_AT: p.R2136* (WGS)
5902_AT: p.N2679fs*9 (WGS)
6018_AT: p.Trp57* (WGS)
6018_AT: p.Tyr1124* (WGS)

Fig. S1. Pathogenic ATM variants in A-T cases. Alignment tracks of whole genome sequencing (WGS) reads at the ATM locus showing pathogenic variants in A-T cases.

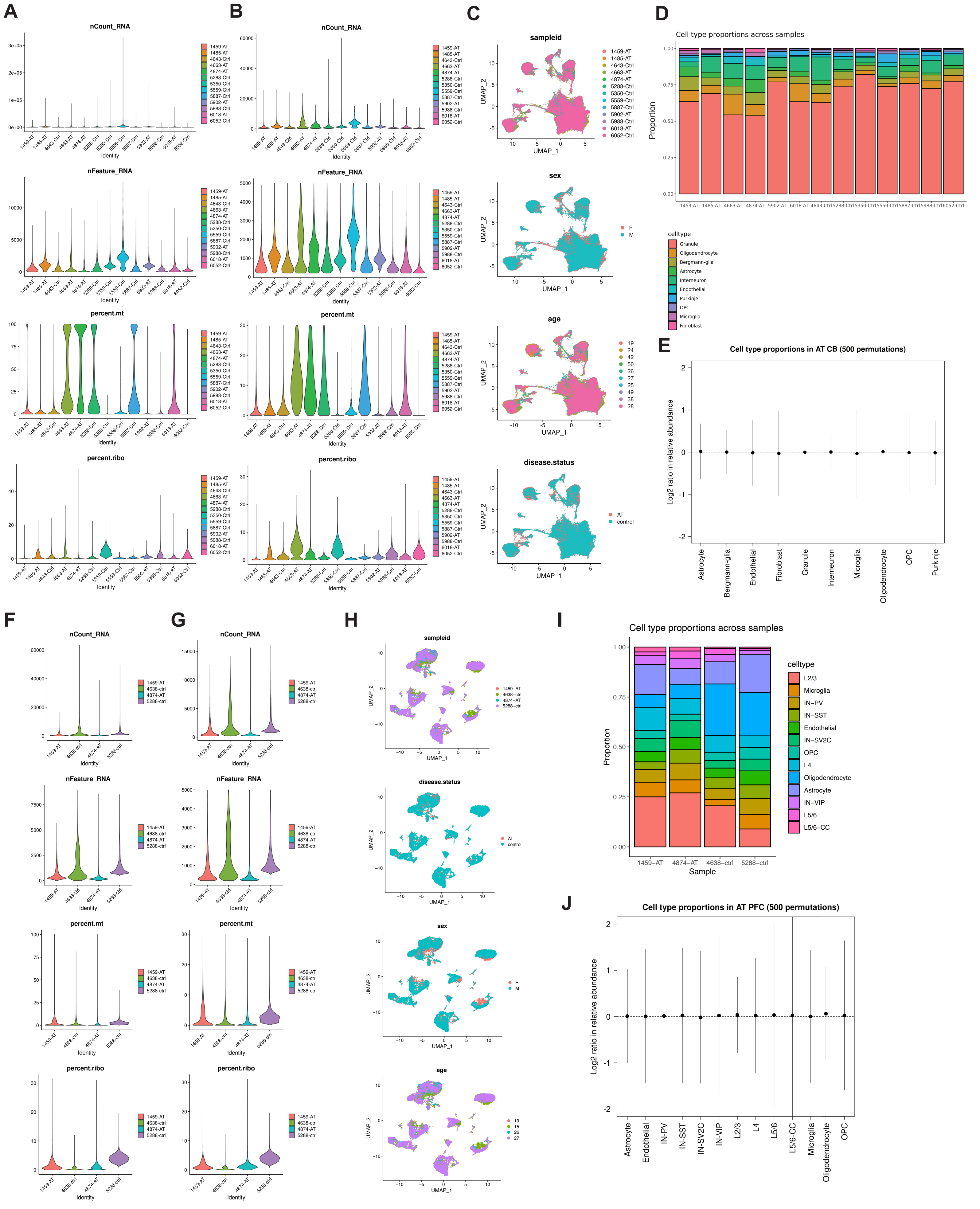

A
B
C
D
nCount_RNA
nFeature_RNA
percent.mt
percent.ribo
Identity
sampleid
sex
age
disease.status
UMAP_1
UMAP_2
Cell type proportions across samples
Proportion
celltype
Granule
Oligodendrocyte
Bergmann-glia
Astrocyte
Interneuron
Endothelial
Purkinje
OPC
Microglia
Fibroblast
E
Cell type proportions in AT CB (500 permutations)
Log2 ratio in relative abundance
F
G
H
I
Sample
L2/3
IN-PV
IN-SST
IN-SV2C
L4
IN-VIP
L5/6
L5/6-CC
J
Cell type proportions in AT PFC (500 permutations)

Fig. S2. Quality control information for snRNA-seq data from cerebellum and PFC. A. Transcript (nCount_RNA), gene (nFeature_RNA), percent transcripts from mitochondrial genes, and percent transcripts from ribosomal genes per cerebellar sample before quality control filtering. B. Transcript (nCount_RNA), gene (nFeature_RNA), percent transcripts from mitochondrial genes, and percent transcripts from ribosomal genes per cerebellar sample after quality control filtering. C. UMAP plots of cerebellar data showing integration of samples, sex, age, and disease status. D. Stacked barplots showing proportion of each cell type per cerebellar sample. E. Posterior distribution of log2 ratio of cell type proportions in A-T cerebellum after 500 random permutations.
F. Transcript (nCount_RNA), gene (nFeature_RNA), percent transcripts from mitochondrial genes, and percent transcripts from ribosomal genes per PFC sample before quality control filtering. G. Transcript (nCount_RNA), gene (nFeature_RNA), percent transcripts from mitochondrial genes, and percent transcripts from ribosomal genes per PFC sample after quality control filtering. H. UMAP plots of cerebellar data showing integration of samples, disease status, sex, and age. I. Stacked barplots showing proportion of each cell type per PFC sample. J. Posterior distribution of log2 ratio of cell type proportions in A-T PFC after 500 random permutations.

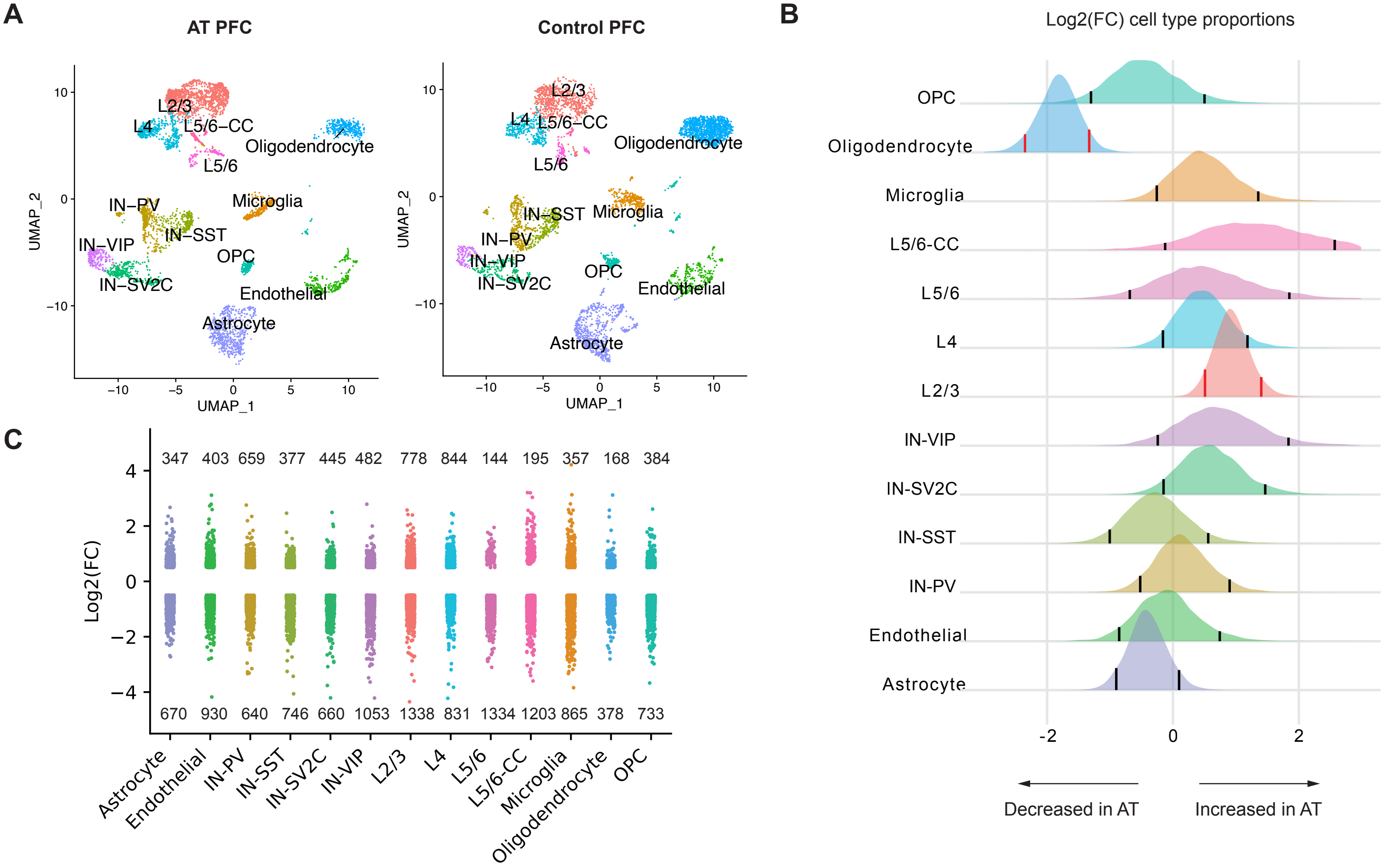


Fig. S3. snRNA-seq data of A-T PFC. A. UMAP plot of major cell types in AT and control human PFC, downsampled to 10,000 cells per condition only for visualization purpose. B. Relative abundance of cell types in AT versus control PFC, shown as the posterior distribution of Log2(proportion in AT/proportion in control) with 89% credible interval. Red bars highlight credible intervals that do not overlap 0. Bolded cell type labels indicate a significant difference in relative abundance. C. Differentially expressed genes (DEGs) in each cell type with FDR<0.05, |Log2FoldChange|>0.50. Each dot represents a significantly differentially expressed gene.

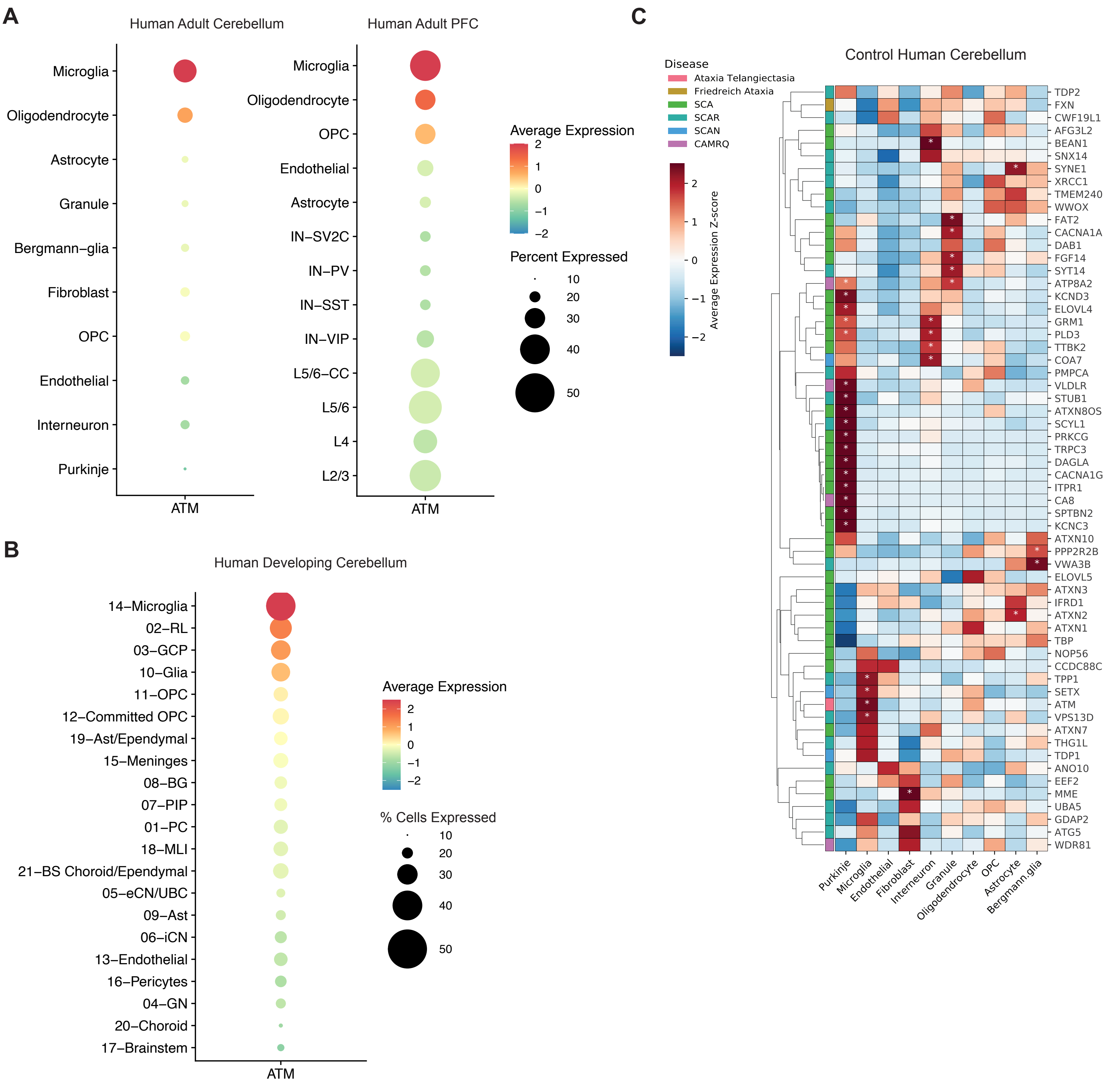


Fig. S4. Enriched expression of monogenic cerebellar disease genes in specific cell types. A. Dotplot of averag

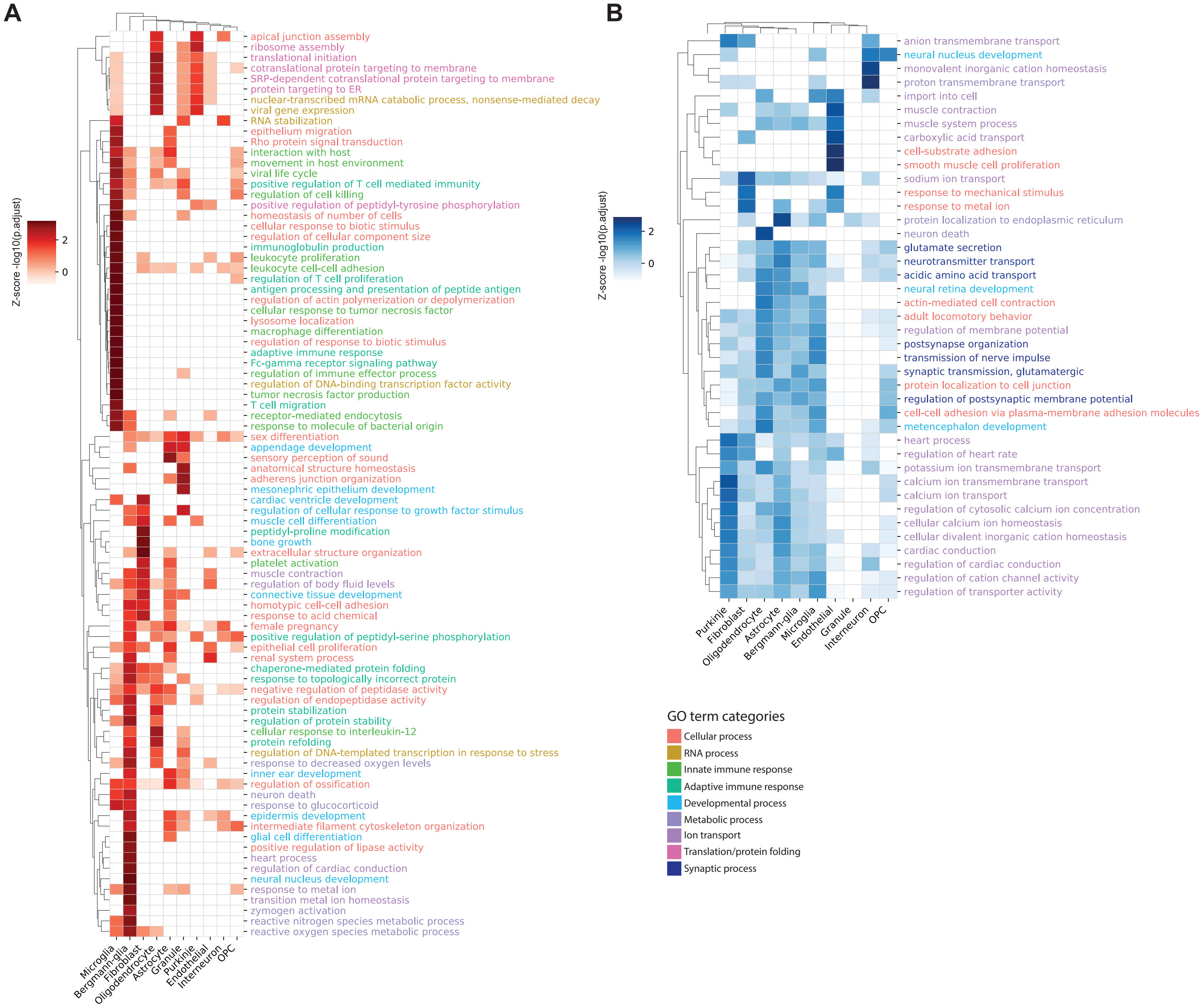


Fig. S5. Gene Ontology enrichment of upregulated and downregulated genes across cell types in A-T cerebellum. A. Heatmap of enriched pathways among upregulated DEGs in cell types from A-T cerebellum. B. Heatmap of enriched pathways among downregulated DEGs in cell types from A-T cerebellum. Color represents z-score of pathway significance (-log10(p. adjusted)) and only significant pathways (p. adj <0.05) are colored.

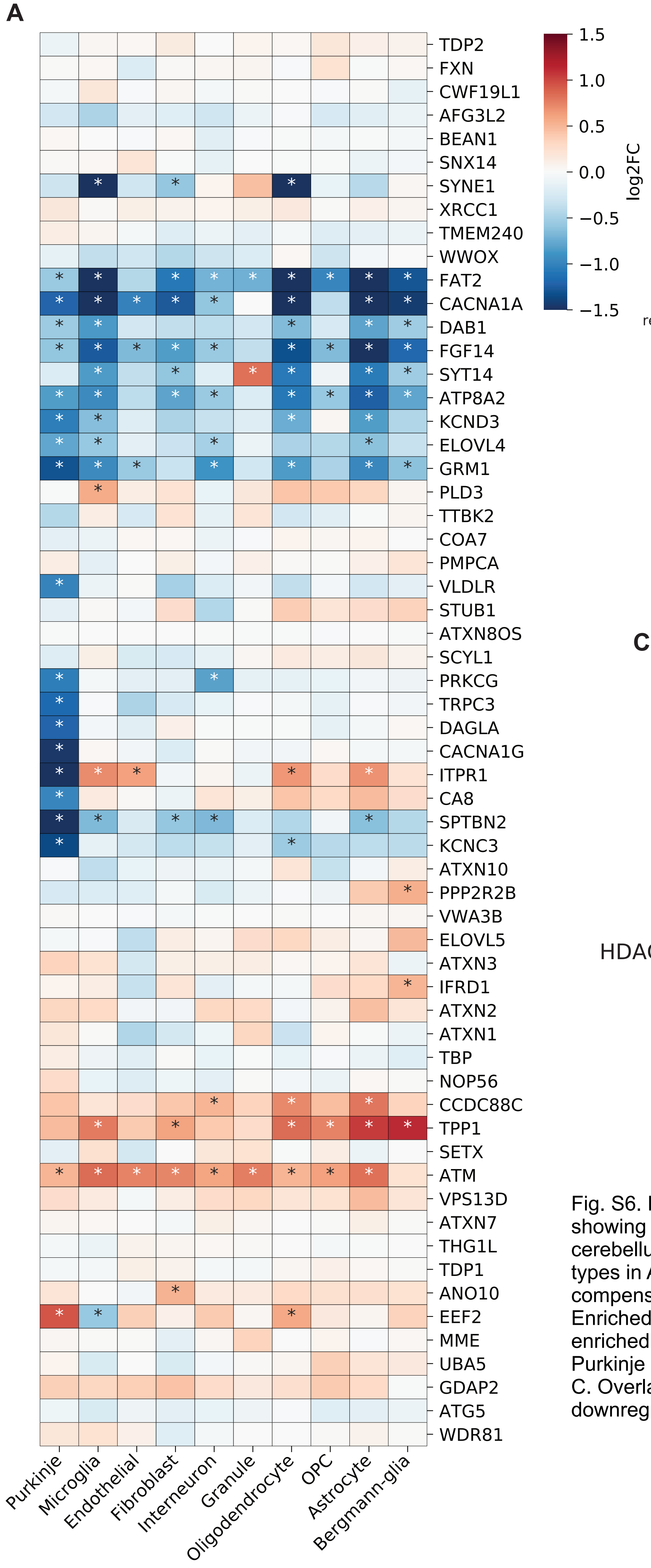


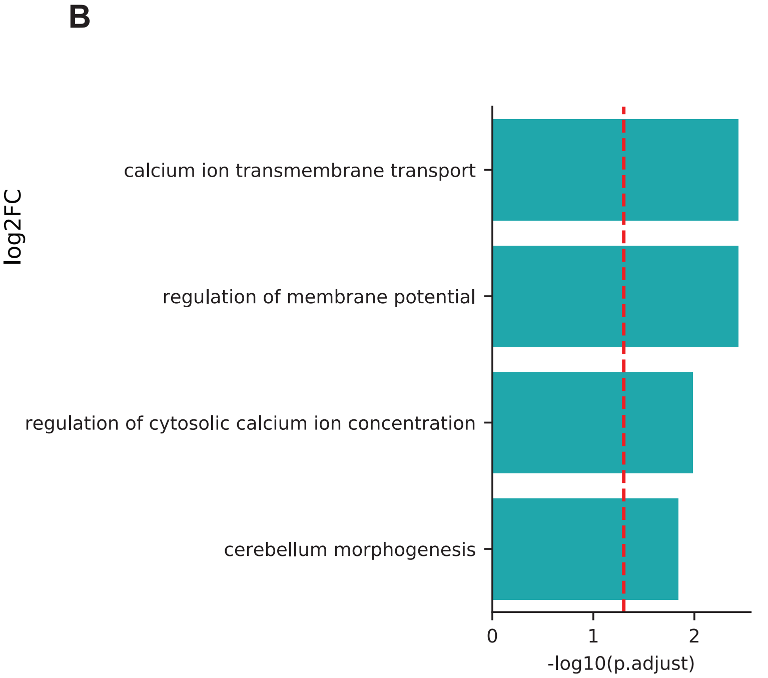


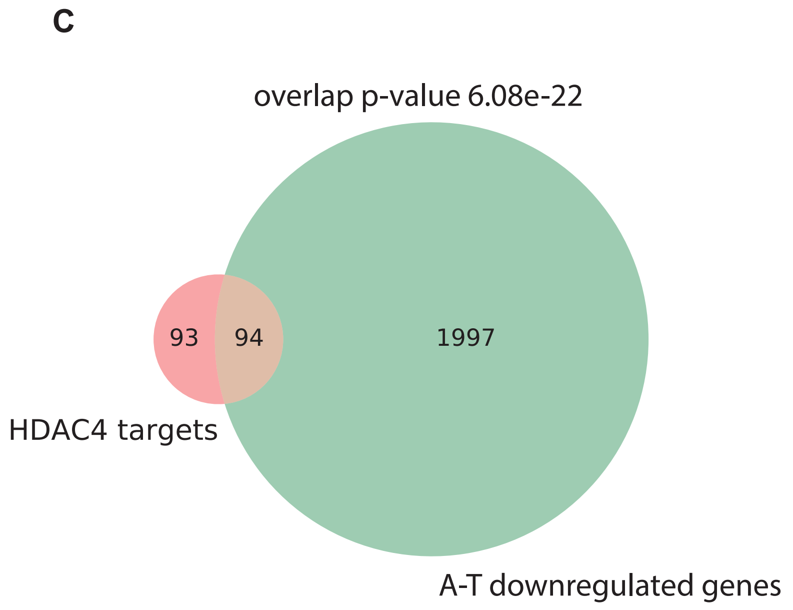


Fig. S6. Dysregulation of hereditary ataxia genes in A-T. A. Heatmap showing log2 fold-change of hereditary ataxia genes in A-T cerebellum. *FDR<0.05. ATM expression increased in several cell types in A-T cerebellum, suggesting that lack of ATM function induces compensatory increases in transcription of the ATM locus. B. Enriched Gene Ontology (GO) pathways among disease genes with enriched expression in Purkinje cells and downregulated in A-T Purkinje cells.
C. Overlap between HDAC4 neuronal target genes and downregulated DEGs in A-T cerebellum.

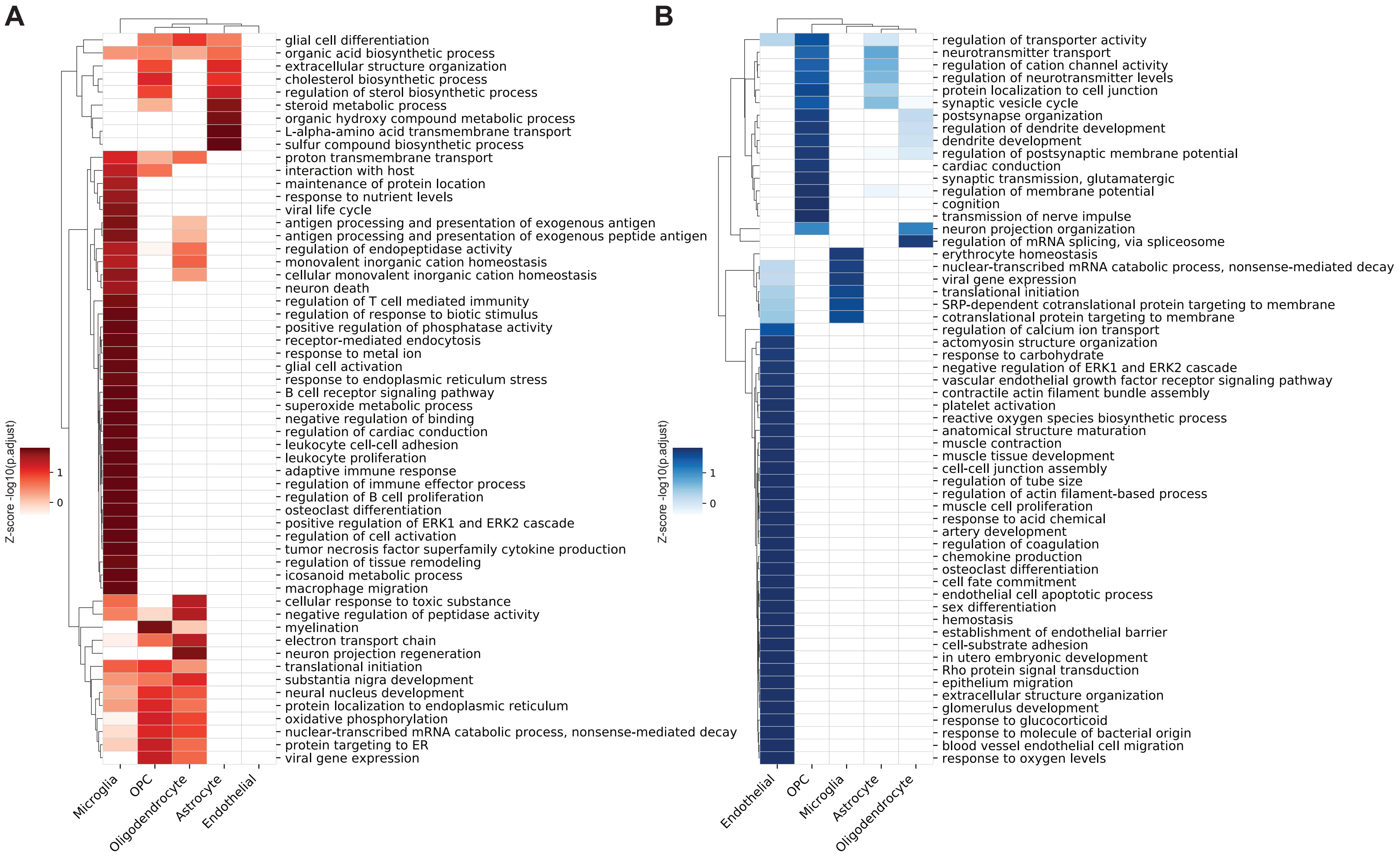


Fig. S7. Enrichment of pathways among genes with greater dysregulation in A-T cerebellum than A-T PFC. A. Heatmap of enriched pathways among DEGs upregulated in AT CB v. AT PFC. B. Heatmap of enriched pathways among DEGs downregulated in AT CB v. AT PFC. Color represents z-score of pathway significance (-log10(p. adjusted)) and only significant pathways (p. adj <0.05) are colored.

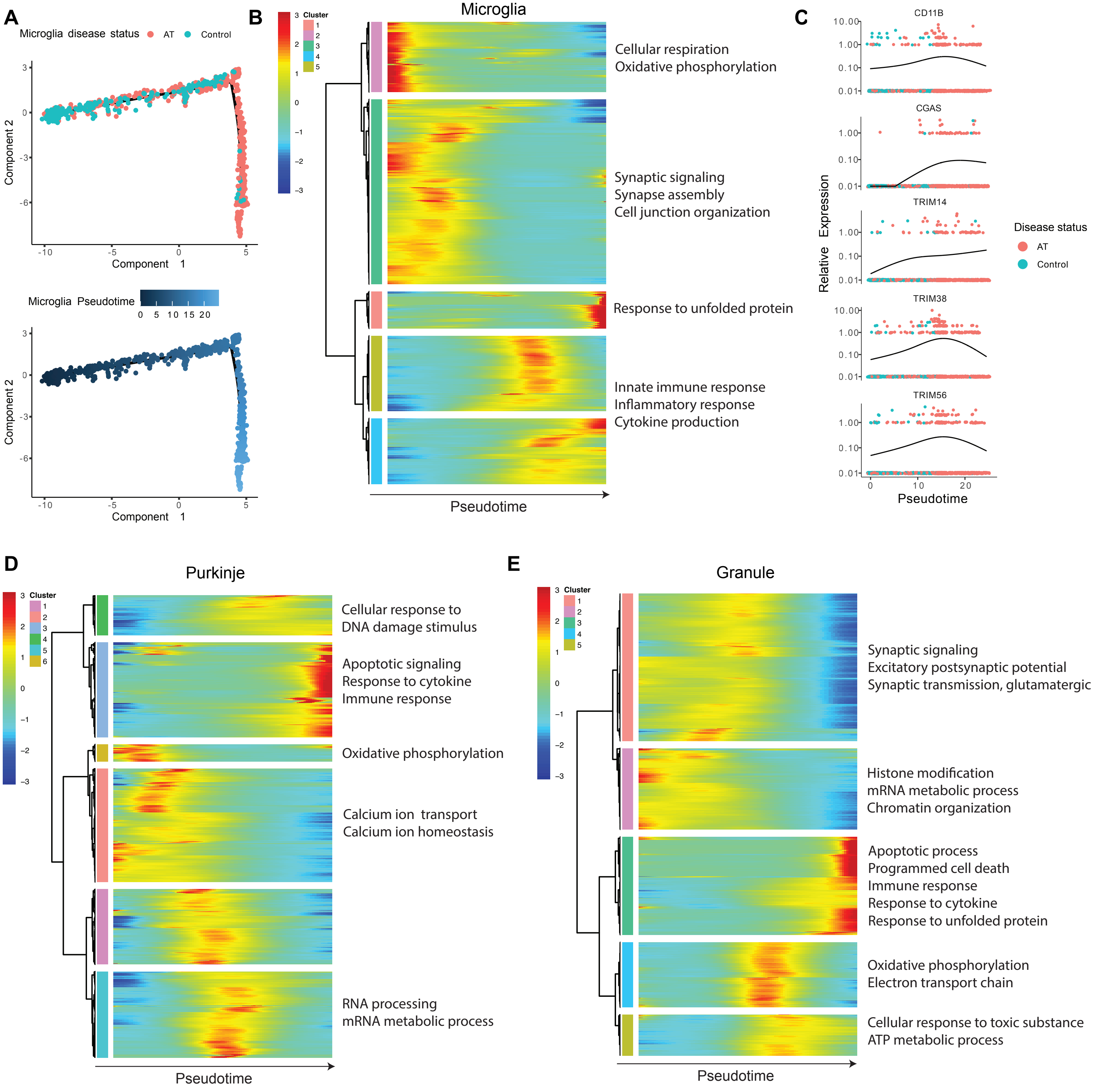


Fig. S8. Pseudotime analysis of disease progression reveals early microglia activation. A. Disease progression pseudotime trajectory of A-T cerebellar microglia, colored by disease status (red: A-T, blue: control), or pseudotime (healthy to diseased). B. Heatmap of genes that significantly change over pseudotime in microglia, clustered by pseudotemporal expression patterns. Each cluster is annotated with enriched GO terms (FDR<0.05). C. Expression of CD11B, CGAS, TRIM14, TRIM38, and TRIM56 over pseudotime in microglia. D. Heatmap of genes that change over pseudotime in Purkinje neurons, clustered by pseudotemporal expression patterns. Each cluster is annotated with enriched GO terms (FDR<0.05). E. Heatmap of genes that change over pseudotime in granule neurons, clustered by pseudotemporal expression patterns. Each cluster is annotated with enriched GO terms (FDR<0.05). Heatmaps in B, D, E depict centered and scaled expression.

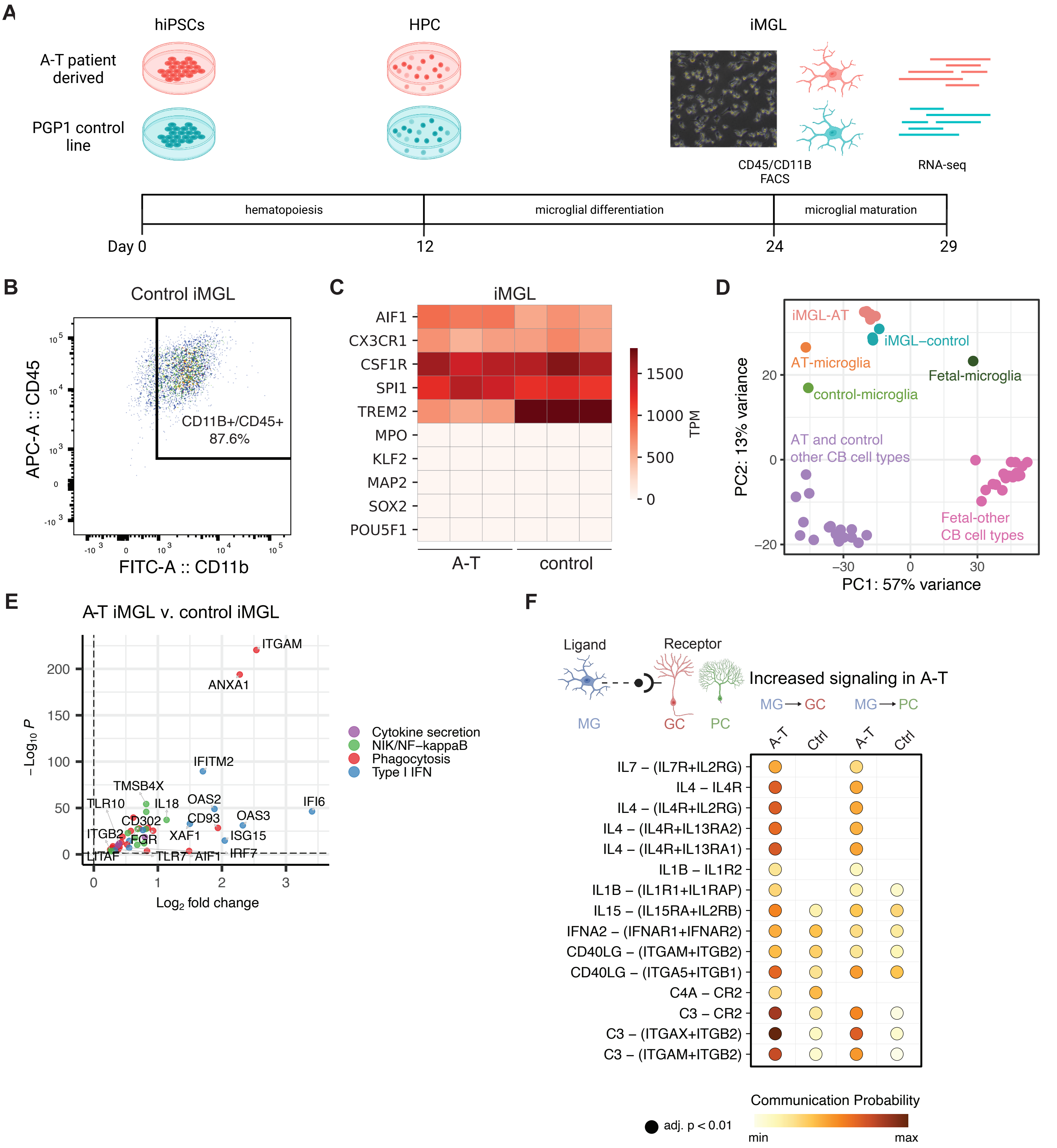
A
hiPSCs
HPC
iMGL
A-T patient derived
PGP1 control line
CD45/CD11B FACS
RNA-seq
hematopoiesis
microglial differentiation
microglial maturation
Day 0
12
24
29
B
Control iMGL
APC-A :: CD45
FITC-A :: CD11b
CD11B+/CD45+ 87.6%
C
iMGL
AIF1
CX3CR1
CSF1R
SPI1
TREM2
MPO
KLF2
MAP2
SOX2
POU5F1
A-T
control
TPM
1500
1000
500
0
D
iMGL-AT
iMGL–control
AT-microglia
Fetal-microglia
control-microglia
AT and control other CB cell types
Fetal-other CB cell types
PC2: 13% variance
PC1: 57% variance
E
A-T iMGL v. control iMGL
ITGAM
ANXA1
IFITM2
TMSB4X
TLR10
IL18
OAS2
IFI6
CD302
CD93
OAS3
ITGB2
FGR
XAF1
ISG15
LITAF
TLR7
AIF1
IRF7
Log2 fold change
Cytokine secretion
NIK/NF–kappaB
Phagocytosis
Type I IFN
F
Ligand
Receptor
MG
GC
PC
Increased signaling in A-T
MG → GC
MG → PC
A-T
Ctrl
IL7 – (IL7R+IL2RG)
IL4 – IL4R
IL4 – (IL4R+IL2RG)
IL4 – (IL4R+IL13RA2)
IL4 – (IL4R+IL13RA1)
IL1B – IL1R2
IL1B – (IL1R1+IL1RAP)
IL15 – (IL15RA+IL2RB)
IFNA2 – (IFNAR1+IFNAR2)
CD40LG – (ITGAM+ITGB2)
CD40LG – (ITGA5+ITGB1)
C4A – CR2
C3 – CR2
C3 – (ITGAX+ITGB2)
C3 – (ITGAM+ITGB2)
Communication Probability
adj. p < 0.01
min
max

Fig. S9. A-T patient iPSC-derived microglia reveal cell-intrinsic activation of NF-kappaB and type I interferon pathways. A. Schematic for generation of iPSC-derived microglia (iMGL) from human A-T patient and control iPSCs. PGP1, Personal Genome Project 1; HPC: hematopoietic progenitor cells. B. Flow cytometry analysis of CD45 and CD11B co-expression in control iMGLs 24 days post-differentiation. C. Expression of microglia (AIF1, CX3CR1, CSF1R, SPI1, TREM2), myeloid lineage (MPO, KLF2), neuronal (MAP2, SOX2), and iPSC (POU5F1) marker genes in A-T and control iMGLs. Each column represents data from iMGL differentiated in an independent well. D. Principal component analysis plot of A-T and control iMGL (iMGL-AT/control) and adult cerebellar cell type pseudobulk transcriptomic profiles derived from snRNA-seq in this study (AT/control-microglia) and fetal cerebellar cell type pseudobulk transcriptomic profiles derived from snRNA-seq data in Aldinger et al., 2021 (Fetal-microglia). E. Volcano plot for log2 fold change in gene expression in A-T versus control iMGL. Representative genes associated with cytokine secretion (purple), NIK/NF-kappaB (green), phagocytosis (red), and type I interferon (IFN) (blue) are highlighted. Horizontal dash line represents FDR=0.05. F. Ligand-receptor pairs with increased communication probability from microglia to granule or Purkinje neurons in A-T cerebellum versus control. Dot color represents communication probability (strength of signaling). Dot size represents significance of ligand-receptor pair and pairs with adjusted p-value <0.01 shown.

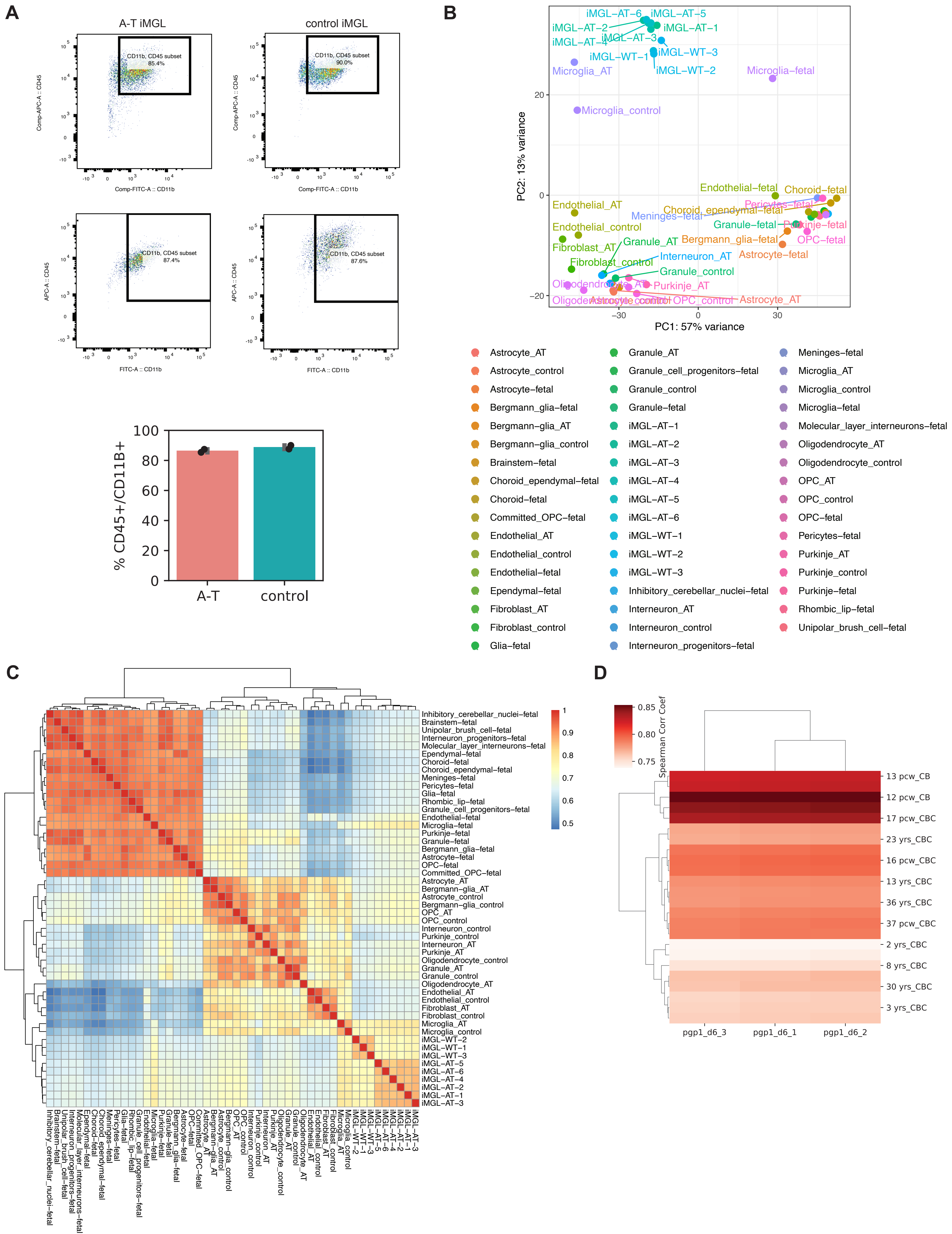
A
A-T iMGL
control iMGL
CD11b, CD45 subset
85.4%
90.0%
87.4%
87.6%
Comp-APC-A :: CD45
Comp-FITC-A :: CD11b
APC-A :: CD45
FITC-A :: CD11b
% CD45+/CD11B+
A-T
control
B
PC1: 57% variance
PC2: 13% variance
iMGL−AT−1
iMGL−AT−2
iMGL−AT−3
iMGL−AT−4
iMGL−AT−5
iMGL−AT−6
iMGL−WT−1
iMGL−WT−2
iMGL−WT−3
Microglia_AT
Microglia_control
Microglia−fetal
Endothelial−fetal
Choroid−fetal
Endothelial_AT
Endothelial_control
Choroid_ependymal−fetal
Pericytes−fetal
Meninges−fetal
Granule−fetal
Purkinje−fetal
OPC−fetal
Bergmann_glia−fetal
Fibroblast_AT
Granule_AT
Interneuron_AT
Fibroblast_control
Astrocyte−fetal
Granule_control
Purkinje_AT
OPC_control
Astrocyte_AT
Astrocyte_AT
Astrocyte_control
Astrocyte−fetal
Bergmann_glia−fetal
Bergmann−glia_AT
Bergmann−glia_control
Brainstem−fetal
Choroid_ependymal−fetal
Choroid−fetal
Committed_OPC−fetal
Endothelial_AT
Endothelial_control
Endothelial−fetal
Ependymal−fetal
Fibroblast_AT
Fibroblast_control
Glia−fetal
Granule_AT
Granule_cell_progenitors−fetal
Granule_control
Granule−fetal
iMGL−AT−1
iMGL−AT−2
iMGL−AT−3
iMGL−AT−4
iMGL−AT−5
iMGL−AT−6
iMGL−WT−1
iMGL−WT−2
iMGL−WT−3
Inhibitory_cerebellar_nuclei−fetal
Interneuron_AT
Interneuron_control
Interneuron_progenitors−fetal
Meninges−fetal
Microglia_AT
Microglia_control
Microglia−fetal
Molecular_layer_interneurons−fetal
Oligodendrocyte_AT
Oligodendrocyte_control
OPC_AT
OPC_control
OPC−fetal
Pericytes−fetal
Purkinje_AT
Purkinje_control
Purkinje−fetal
Rhombic_lip−fetal
Unipolar_brush_cell−fetal
C
Inhibitory_cerebellar_nuclei−fetal
Brainstem−fetal
Unipolar_brush_cell−fetal
Interneuron_progenitors−fetal
Molecular_layer_interneurons−fetal
Ependymal−fetal
Choroid−fetal
Choroid_ependymal−fetal
Meninges−fetal
Pericytes−fetal
Glia−fetal
Rhombic_lip−fetal
Granule_cell_progenitors−fetal
Endothelial−fetal
Microglia−fetal
Purkinje−fetal
Granule−fetal
Bergmann_glia−fetal
Astrocyte−fetal
OPC−fetal
Committed_OPC−fetal
Astrocyte_AT
Bergmann−glia_AT
Astrocyte_control
Bergmann−glia_control
OPC_AT
OPC_control
Interneuron_control
Purkinje_control
Interneuron_AT
Purkinje_AT
Oligodendrocyte_control
Granule_AT
Granule_control
Oligodendrocyte_AT
Endothelial_AT
Endothelial_control
Fibroblast_AT
Fibroblast_control
Microglia_AT
Microglia_control
iMGL−WT−2
iMGL−WT−1
iMGL−WT−3
iMGL−AT−5
iMGL−AT−6
iMGL−AT−4
iMGL−AT−2
iMGL−AT−1
iMGL−AT−3
D
Spearman Corr Coef
13 pcw_CB
12 pcw_CB
17 pcw_CBC
23 yrs_CBC
16 pcw_CBC
13 yrs_CBC
36 yrs_CBC
37 pcw_CBC
2 yrs_CBC
8 yrs_CBC
30 yrs_CBC
3 yrs_CBC
pgp1_d6_3
pgp1_d6_1
pgp1_d6_2

Fig. S10. Differentiation of A-T patient and control human iPSCs into microglia and neurons. A. Flow cytometry analysis of CD45 and CD11B co-expression in A-T and control iMGLs 24 days post-differentiation from 2 independent differentiation batches. B. Principal component analysis plot of A-T and control iMGL and adult cerebellar cell type pseudobulk transcriptomic profiles derived from snRNA-seq in this study and fetal cerebellar cell type pseudobulk transcriptomic profiles derived from snRNA-seq data in Aldinger et al., 2021. C. Heatmap of Spearman Correlation Coefficients between human adult and fetal cerebellar cell type transcriptome profiles and iMGL transcriptome profiles. D. Heatmap of Spearman Correlation Coefficients between human control (PGP1) iPSC-derived neurons at day 6 of differentiation transcriptomes and human developing cerebellum (CB) and cerebellar cortex (CBC) transcriptomes from BrainSpan: Atlas of the Developing Human Brain. iMGL-AT, A-T iPSC-derived microglia. iMGL-WT, control iPSC-derived microglia.

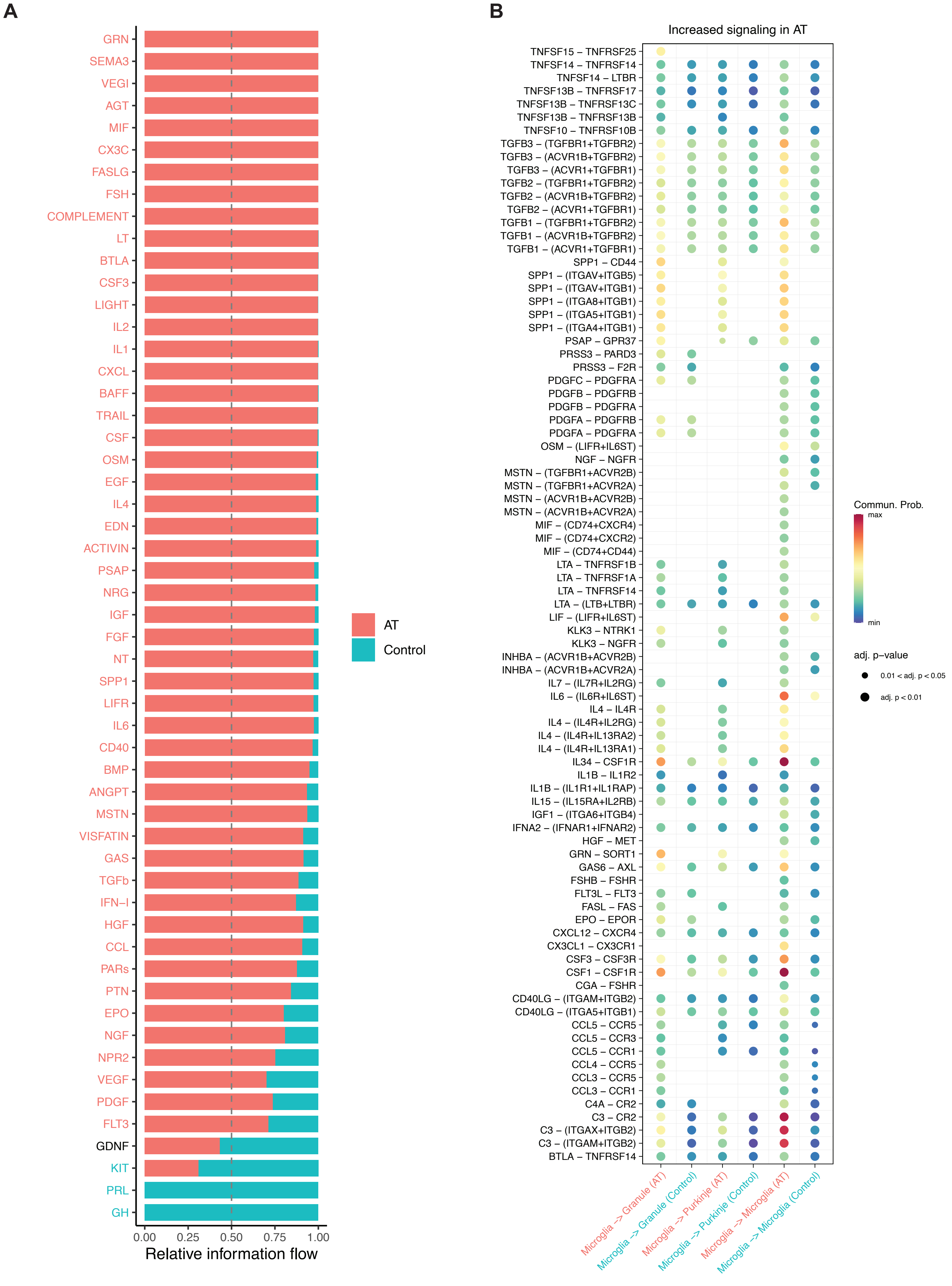

A
GRN
SEMA3
VEGI
AGT
MIF
CX3C
FASLG
FSH
COMPLEMENT
LT
BTLA
CSF3
LIGHT
IL2
IL1
CXCL
BAFF
TRAIL
CSF
OSM
EGF
IL4
EDN
ACTIVIN
PSAP
NRG
IGF
FGF
NT
SPP1
LIFR
IL6
CD40
BMP
ANGPT
MSTN
VISFATIN
GAS
TGFb
IFN−I
HGF
CCL
PARs
PTN
EPO
NGF
NPR2
VEGF
PDGF
FLT3
GDNF
KIT
PRL
GH
0.00
0.25
0.50
0.75
1.00
Relative information flow
AT
Control
B
Increased signaling in AT
TNFSF15 – TNFRSF25
TNFSF14 – TNFRSF14
TNFSF14 – LTBR
TNFSF13B – TNFRSF17
TNFSF13B – TNFRSF13C
TNFSF13B – TNFRSF13B
TNFSF10 – TNFRSF10B
TGFB3 – (TGFBR1+TGFBR2)
TGFB3 – (ACVR1B+TGFBR2)
TGFB3 – (ACVR1+TGFBR1)
TGFB2 – (TGFBR1+TGFBR2)
TGFB2 – (ACVR1B+TGFBR2)
TGFB2 – (ACVR1+TGFBR1)
TGFB1 – (TGFBR1+TGFBR2)
TGFB1 – (ACVR1B+TGFBR2)
TGFB1 – (ACVR1+TGFBR1)
SPP1 – CD44
SPP1 – (ITGAV+ITGB5)
SPP1 – (ITGAV+ITGB1)
SPP1 – (ITGA8+ITGB1)
SPP1 – (ITGA5+ITGB1)
SPP1 – (ITGA4+ITGB1)
PSAP – GPR37
PRSS3 – PARD3
PRSS3 – F2R
PDGFC – PDGFRA
PDGFB – PDGFRB
PDGFB – PDGFRA
PDGFA – PDGFRB
PDGFA – PDGFRA
OSM – (LIFR+IL6ST)
NGF – NGFR
MSTN – (TGFBR1+ACVR2B)
MSTN – (TGFBR1+ACVR2A)
MSTN – (ACVR1B+ACVR2B)
MSTN – (ACVR1B+ACVR2A)
MIF – (CD74+CXCR4)
MIF – (CD74+CXCR2)
MIF – (CD74+CD44)
LTA – TNFRSF1B
LTA – TNFRSF1A
LTA – TNFRSF14
LTA – (LTB+LTBR)
LIF – (LIFR+IL6ST)
KLK3 – NTRK1
KLK3 – NGFR
INHBA – (ACVR1B+ACVR2B)
INHBA – (ACVR1B+ACVR2A)
IL7 – (IL7R+IL2RG)
IL6 – (IL6R+IL6ST)
IL4 – IL4R
IL4 – (IL4R+IL2RG)
IL4 – (IL4R+IL13RA2)
IL4 – (IL4R+IL13RA1)
IL34 – CSF1R
IL1B – IL1R2
IL1B – (IL1R1+IL1RAP)
IL15 – (IL15RA+IL2RB)
IGF1 – (ITGA6+ITGB4)
IFNA2 – (IFNAR1+IFNAR2)
HGF – MET
GRN – SORT1
GAS6 – AXL
FSHB – FSHR
FLT3L – FLT3
FASL – FAS
EPO – EPOR
CXCL12 – CXCR4
CX3CL1 – CX3CR1
CSF3 – CSF3R
CSF1 – CSF1R
CGA – FSHR
CD40LG – (ITGAM+ITGB2)
CD40LG – (ITGA5+ITGB1)
CCL5 – CCR5
CCL5 – CCR3
CCL5 – CCR1
CCL4 – CCR5
CCL3 – CCR5
CCL3 – CCR1
C4A – CR2
C3 – CR2
C3 – (ITGAX+ITGB2)
C3 – (ITGAM+ITGB2)
BTLA – TNFRSF14
Microglia -> Granule (AT)
Microglia -> Granule (Control)
Microglia -> Purkinje (AT)
Microglia -> Purkinje (Control)
Microglia -> Microglia (AT)
Microglia -> Microglia (Control)
Commun. Prob.
max
min
adj. p−value
0.01 < adj. p < 0.05
adj. p < 0.01

Fig. S11. Altered cell-cell communication in A-T cerebellum. A. Relative information flow for signaling pathways enriched in A-T or control cerebellum. Information flow for each pathway is calculated as the sum of the communication probability among all pairs of cell groups. B. Ligand-receptor pairs with increased communication probability between microglia, granule, and Purkinje neurons in A-T cerebellum compared to control. Dot color represents communication probability (strength of signaling) and dot size represents significance of ligand-receptor pair (adjusted p-value). Empty space represents a communication probability of zero.
